# Differential modulation of aversive signaling by expectation across the cingulate cortex

**DOI:** 10.64898/2026.01.05.697709

**Authors:** Sophie A. Rogers, Corinna S. Oswell, Lindsay L. Ejoh, Gregory Corder

## Abstract

Pain-related aversion is an affective-motivational state driven by sensory experience that promotes learning and recruits widespread cortical networks, yet how distinct cingulate subregions contribute to its adaptive utility remains poorly understood. Here we used longitudinal one-photon calcium imaging in mice to compare dynamics in the anterior cingulate cortex (ACC) and retrosplenial cortex (RSC) across repeated unsignaled foot-shocks and fear conditioning and extinction paradigm. Both regions contained relatively stable ensembles that responded robustly to shocks, indicating shared encoding of acute nociceptive events. However, only the RSC flexibly re-organized its population activity when shocks were preceded by predictive cues. These anticipatory dynamics in the RSC predicted the rate of fear learning across individuals and subsequent extinction. By contrast, the ACC maintained shock-responsive ensembles with limited cue modulation. Instead, its dynamics encoded decisions to freeze, aligning with its role in encoding ongoing nociception and driving immediate defensive behavior. Together, these results reveal a division of labor in which the ACC emphasizes ongoing nociceptive processing, while the RSC transforms sensory signals into predictive codes that shape learning and memory. This specialization highlights how distributed cortical computations cooperate to generate the adaptive value of aversion. More broadly, our findings suggest that these regions assume complementary roles to address immediate sensory-motivational responses while flexibly reconfiguring to support long-term behavioral adaptation.

**Significance statement:** Pain engages widespread cortical circuits, yet how distinct cingulate subregions collaborate to shape its experience and utility remains unknown. Using longitudinal calcium imaging in mice, we demonstrate that both the anterior cingulate and retrosplenial cortex contain stable shock-responsive ensembles, but only the retrosplenial cortex flexibly remodels its activity when shocks are predicted by cues. These anticipatory dynamics not only predict fear learning but influence extinction. Our findings uncover a division of labor in which the anterior cingulate encodes ongoing nociception and immediate defensive actions, while the retrosplenial cortex transforms these signals into temporally structured representations that support learning and memory. This work highlights how specialized cortical computations interact to generate the adaptive value of pain.

## Introduction

Aversive experiences generated by noxious stimuli trigger a rapid, prepotent cascade of sensation, perception, motivation, action, and learning, engaging the entire brain^1,2^. The cingulate cortex, an anatomical continuum consisting of the anterior cingulate and retrosplenial cortex (ACC and RSC), comprises the crucial cortical hub for pain and fear^3–8^. These regions’ overlapping involvement in pain-related aversion is at odds with their otherwise disparate functions: the ACC in decision-making and behavioral selection and the RSC in spatial cognition and episodic memory^9–15^. How their distinct specializations define their roles in aversive processing is unknown.

Both regions are robustly activated in humans and rodents in response to noxious stimuli^16–19^. Both undergo robust consequent transcriptional and synaptic plasticity^4,20,21^, and pharmacological, optical, and chemogenetic inhibition of neural ensembles in both regions can produce analgesia^22–27^. We have previously shown that ensembles in the ACC are involved in recuperative behaviors following acute and chronic injury^25^. However, despite the presence of stable, nociception-discriminating population dynamics, the presence of a stable nociceptive ensemble the ACC is hotly debated and has yet to be investigated in the RSC^25,28^.

Another function both regions share is the mediation of aversive learning; in particular, directing learning about noxious stimuli. In one paper, glutamatergic activation of the ACC was necessary and sufficient for conditioned place aversion, leading the authors to conclude that the ACC produces an “aversive teaching signal”^29^. Another group found that optogenetic reactivation of FOS-expressing neurons that were active during contextual fear conditioning in the RSC was sufficient to reinstate fear following extinction^30^.

One form of aversive learning for which both regions are crucial is trace fear conditioning (TFC). In TFC, a foot-shock follows a conditioned cue by a window of 10-30 seconds called the trace period. Glutamatergic plasticity in the ACC, which is linked to nociception, is necessary and sufficient for TFC acquisition^31–35^. In particular, neurons in the ACC become tuned to freezing-on or freezing-off bouts over fear conditioning^36,37^. The same is required not only for acquisition, but also for extinction in the RSC^38–44^. Indeed, we previously showed that the persistent activation of neurons recruited during TFC acquisition over extinction predicted enhanced fear recall three days later in mice^45^.

To explore how these functionally disparate but anatomically connected regions participate in aversive sensation, behavior, and learning, we performed longitudinal one-photon calcium imaging in mice implanted with GRIN lenses in either the ACC or the RSC in two experiments. First, in our “unsignaled shock” experiment, mice underwent repeated foot-shocks minutes, hours, days, and weeks apart to determine how stable single cell responses and population dynamics were over time. Second, we performed the TFC protocol described in our previous work^45^ in separate cohorts of mice to examine how these dynamics might be modulated by expectation of shock and how they might influence learning.

Given the ACC’s well-established role in nociception and negative affect, we hypothesized that the ACC would contain a stable shock-responsive population over days and that their activity would be predictive of fear recall. Given the RSC’s role in spatio-temporal computation and memory, we hypothesized that it would lack stable dynamics to unsignaled shock, but that it would recruit and modulate responses in neural ensembles over TFC acquisition and extinction that would be predictive of learning. Instead, we found that both regions contained relatively stable neural and population-level dynamics over time in the unsignaled shock experiment, with subtle and interesting distinctions, but that only activity in the RSC was enhanced by the expectation of an upcoming shock during TFC acquisition and predictive of both fear recall and extinction. Together, these findings improve our understanding of how functional specializations allow these inter-connected regions to collaborate to produce the experience and utility of aversion.

## Results

### Cingulate neurons encode shock stably within a day, drifts over weeks

First, we aimed to investigate whether there are stable neural ensembles, single cell dynamics, and population dynamics in response to repeated foot-shocks across varying timescales in each region. Mice were placed in a Med Associates chamber with an electrified grid floor over five sessions, 4 hours, 24 hours, 1 week, and 2 weeks apart from the first session (0hours) (**Fig. 1A**). Eight footshocks (1mA, 2sec) were delivered at pseudorandomized jittered intervals 60±10sec apart from one another. We imaged and longitudinally registered 596 neurons from four mice expressing AAV9-*Syn*-jGaMP8m and implanted with 1-mm diameter Inscopix GRIN lenses over the ACC and 147 neurons from three mice implanted over RSC (**Fig. 1B,C, S1A**).

**Figure 1.**
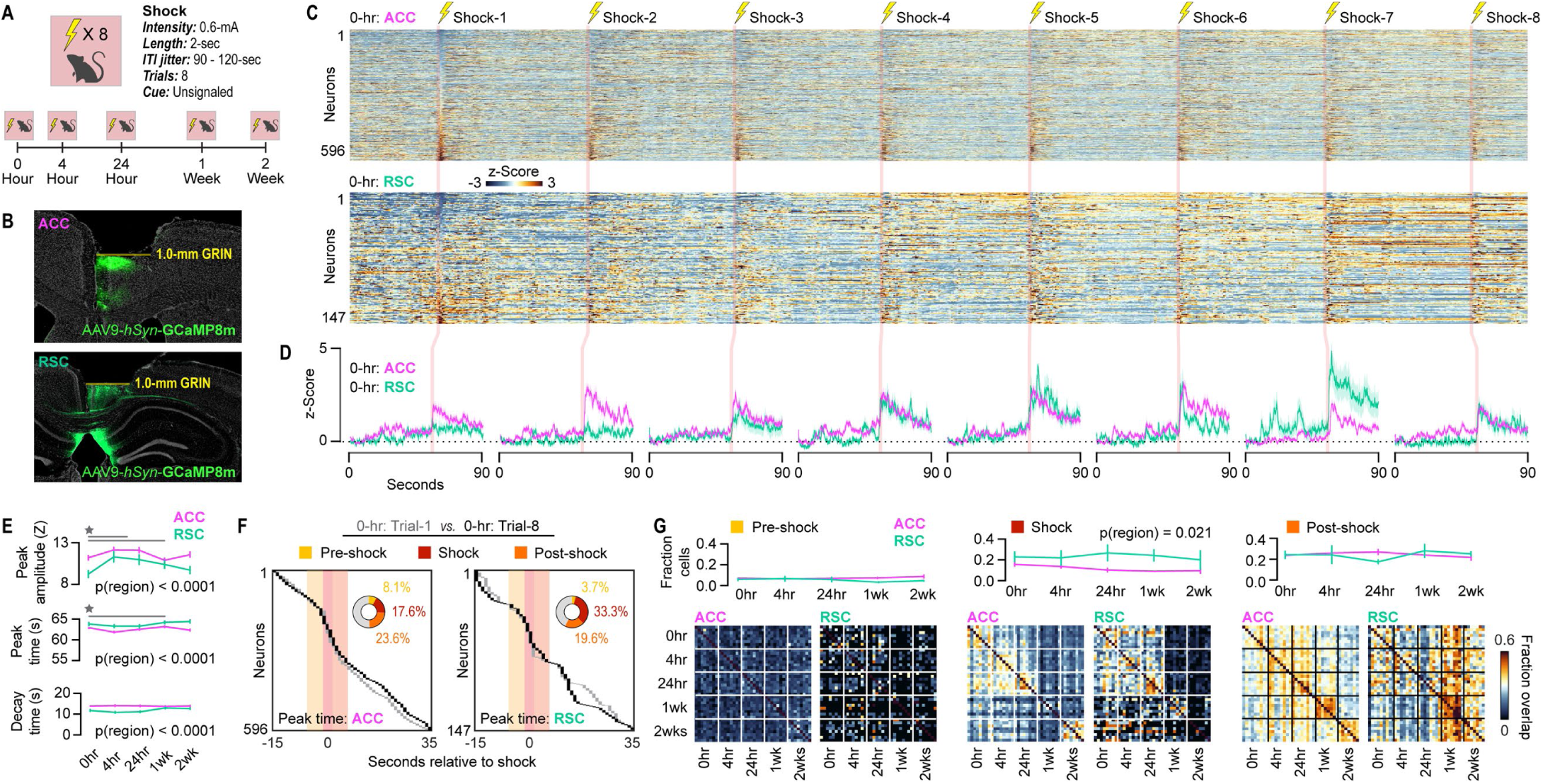
ACC and RSC neurons encode unsignaled shock. **A.** Experimental diagram. **B.** Left: image of implantation area over ACC, with GCaMP8m-expressing neurons in green and DAPI in white. Right: Heatmap of longitudinally registered ACC neuron activity (z-score) aligned to shocks at 0hours. **C.** Left: image of implantation area over RSC. Right: Heatmap of longitudinally registered RSC neuron activity (z-score) aligned to shocks at 0hours. **D.** Average and SEM PSTHs aligned to shock delivery over trials 1-8 at 0hours in the ACC (pink, n=596) and RSC (green, n=147). **E.** Across all ACC and RSC neurons, average peak amplitude of signal (left; Two-Way ANOVA: pSession = 0.0043, F(4,3705) = 3.81; pRegion < 0.0001, F(1,3705) = 17.87), average peak time (middle; Two-Way ANOVA: pSession = 0.0043, F(4,3705) = 3.82; pRegion < 0.0001, F(1,3705) = 38.27), and average decay time (right; Two-Way ANOVA: pRegion < 0.0001, F(1,3705) = 41.47). Red bars indicate shock delivery. **F.** Left: time of peak signal for each ACC neuron during trial 1 (faded) and trial 8 (solid) of Acquisition. Inset: proportion of neurons peaking within 5s prior to shock (Pre-shock neurons, yellow), during shock (Shock neurons, red), and within 5s post-shock, (Post-shock neurons, orange). Right: Same for RSC neurons. (Chi Square test between regions: p = 0.11, Chi-Square(3) = 7.30). **G.** Left: Fraction of Pre-shock neurons across animals (top; nACC = 4, nRSC = 3; Two-Way RM ANOVA) and fraction overlap of Pre-shock neurons between each trials (bottom), with white dashed lines indicating new session, in ACC (left) and RSC (right). Middle: Same as Top for Shock neurons (pRegion = 0.022, F(1,5) = 10.76). Right: Same is H for Post-shock neurons. Stars indicate p<0.05. Green stars indicate change in RSC over time; pink stars indicate change in ACC over time; gray stars with lines indicate change across regions over time; gray stars without lines indicate difference between regions. Statistics in Supplementary Table Rows 1-7.

Just under half of neurons in each region were significantly up- or down-regulated to shock on any given trial (**Fig. S1B; Supp. Table Rows 86-95**). Over trials in the first session, the fraction of shock-upregulated RSC neurons increased, while the fraction of downregulated neurons decreased, indicating some population-level sensitization to unsignaled shock over minutes (**Fig. S1B**). In both regions, neurons that were shock up-regulated on one trial were upregulated on 40-45% of all other trials across sessions, while neurons that were shock down-regulated were much less stable, particularly in the RSC (**Fig. S1C, D; Supp. Table Row 96**). We then dissected the dynamics of these responses more granularly. In the first two trials of the first session, foot-shocks only increased ACC activity, with signal decaying on the order of 10s of seconds (**Fig. 1D**). RSC populations gradually acquired similar dynamics, with increased activation in response to shock over trials. Averaging over trials from the same session, RSC neurons consistently peaked with lower amplitude, at later times, and with faster decay times than ACC neurons (**Fig. 1E, S1E-H, Supp. Table Rows 1-3**). Interestingly, response amplitudes increased in both populations between the first and the second and third sessions (4hours and 24hours), suggesting that repeated exposure to the same aversive stimulus within a day sensitizes cingulate responses. Time to peak also increased between 4hours and 1week, suggesting increased length of time over which neurons display heightened activity. Decay times did not change over sessions and were thus unaffected by re-exposure to shock.

To further investigate the physiological meaning of response peak timing, we identified neurons that peak in the 5 seconds before the shock delivery (Pre-shock neurons), during the 2 second shock itself (Shock neurons), or in the 5 seconds after shock delivery (Post-shock neurons, **Fig. 1F, Supp. Table Row 4**). Proportions of these neurons on the last trial of the first session were similar between regions, with Pre-shock neurons making up the lowest proportion (ACC: 8.1%, n = 49; RSC: 3.7%, n = 5) and Shock (ACC: 17.6%; n = 115, RSC: 33.3%, n = 45) and Post-shock neurons (ACC: 23.6%, n = 127; RSC: 19.6%, n = 33). Across sessions, these proportions did not change (**Fig. 1G, top, Supp. Table Rows 5-7**). Surprisingly, however, the RSC hosted a significantly greater population of Shock neurons than the ACC.

Pre-shock neurons were largely unstable, with little overlap between trials or sessions (mean_ACC between_ = 8.8%, mean_ACC, within_ = 23.3%, mean_RSC between_ = 6.5%, mean_RSC, within_ = 17.9%; **Fig. 1G, bottom left, S2**). In contrast, Shock neurons were highly stable within sessions and between 0-24hours, suggesting that a stable shock-responsive population *does* exist, in *both* regions, but that representational drift occurs on the order of days and weeks (mean_ACC between_ = 25.6%, mean_ACC, within_ = 45.9%, mean_RSC between_ = 23.3%, mean_RSC, within_ = 42.0%; **Fig. 1G, bottom middle, S2**). Only post-shock neuron stability qualitatively differed between regions (mean_ACC between_ = 24.4%, mean_ACC, within_ = 39.3%, mean_RSC between_ = 24.4%, mean_RSC, within_ = 38.0%; **Fig. 1G, bottom right, S2**). Post-shock neurons were highly stable within and between sessions in the ACC, with some drop out occurring by 2weeks. In contrast, Post-shock neurons were highly *unstable* in the RSC – until 1week. Post-shock neurons at 1 and 2weeks in the RSC overlap highly with Post-shock neurons from all earlier trials, suggesting that, as time progresses, the RSC recruits neurons that were active following a salient or aversive stimulus to represent that stimulus weeks later. These results are highly consistent with evidence for a stable role for ACC in nociception and aversion and that for RSC in long-term or remote episodic memory.

### Distinct ensembles drive population coding of shock in each region

In Kanter et al., 2025, the authors found that trajectory accelerations in neural subspaces indicated “boundary”-like events, where, around a salient event, entorhinal cortex trajectories would rapidly jump to a new subspace^46^. These jumps, or changes in speed, can be quantified as acceleration of neural trajectories through a latent space. To compute this, we considered the first two principle components (PCs) of the ACC and RSC populations in each session (**Fig. 2A**). We then calculated the second derivative of the Euclidean coordinates over time in each animals’ PC space and averaged over trials. As expected, we observed trajectory accelerations in both regions at 0hours, 4hours, 24hours, and 1week (**Fig. 2B, Supp. Table Rows 8-12**). Surprisingly, no trajectory acceleration was observed at 2weeks, indicating an effect of either over-familiarization or contextual memory. Perhaps, by the fifth session, the familiar context was sufficient to induce entry into a contextual fear subspace, eliminating the requirement for trajectory accelerations to enter that subspace; alternatively, but not mutually exclusively, these accelerations could indicate prediction errors that were present when the shocks were still surprising, and their absence could indicate the ultimate absence of surprise.

**Figure 2.**
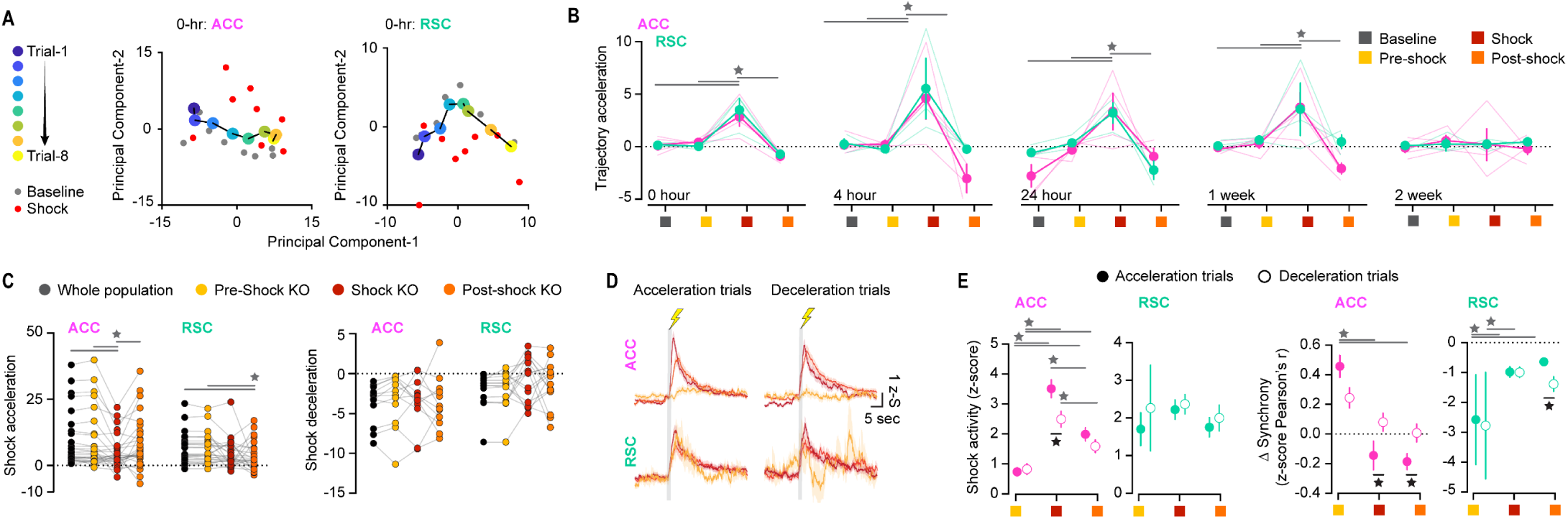
Distinct temporal dynamics accelerate ACC and RSC trajectories during unsignaled shocks. **A.** PCA of ACC (left) and RSC (right) activity over 0hours. Large dots are trial averaged coordinates. Dark blue indicates the first trial and bright yellow indicates the last. Gray dots are averaged coordinates during the baseline periods of each trial. Red dots are averaged coordinates during the shocks. **B.** ACC and RSC trial-averaged trajectory accelerations during each trial period in each session. Dots and error bars are mean and SEM; lines are individual animals. (Two-Way RM ANOVA: pTrialPeriod_0hours_ < 0.0001, F(3,15) = 22.47; pTrialPeriod_0hours_ = 0.0019, F(3,15) = 8.11; pTrialPeriod_0hours_= 0.0016, F(3,15) = 8.48; pTrialPeriod_1week_ = 0.0041, F(3,15) = 6.80; pTrialPeriod_2weeks_ = 0.96, F(3,15) = 0.10). **C.** Left: Comparison of shock-associated trajectories across each condition in trials where shocks successfully evoked an acceleration in the ACC (left, One-Way ANOVA: pCondition < 0.0001, F(3,81) = 8.00) and RSC (right, One-Way ANOVA: pCondition = 0.0143, F(3,72) = 0.014). Right: Same as Left for trials without accelerations in the ACC (left) and RSC (right). **D.** Mean and SEM PSTHs for each population of neurons in the ACC (top) and RSC (bottom) in deceleration (left) and acceleration (right) trials. **L.** Left: activity during the shock in each population of neurons in the ACC (left; pInteraction = 0.050, F(2,636) = 2.99) and RSC (right) during acceleration (solid dot) and deceleration (empty dot) trials. Right: change in synchrony (z-score Pearson’s r) during the shock with respect to baseline in each population of neurons in the ACC (left; pInteraction = 0.031, F(2,636) = 3.47) and RSC (right; pInteraction = 0.048, F(2,160) = 0.048) during acceleration (solid dot) and deceleration (empty dot) trials. Stars indicate p<0.05. Gray stars with lines indicate change over time; black stars indicate difference between groups. Statistics in Supplementary Table Rows 8-20.

To determine whether peak time-defined ensembles of neurons were important for these trajectory accelerations, this procedure was repeated on neurons pooled across animals in each region and again after serially removing each subpopulation from analysis. Intriguingly, different ensembles were necessary for these shock-evoked trajectory accelerations. Shock-evoked trajectory accelerations were dependent on Shock neurons in ACC and Post-shock neurons in RSC (**Fig. 2C left, Supp. Table Rows 13-14**). Although the RSC contained more Shock neurons than ACC, they are inessential for shock-evoked population dynamics, showing that this effect is independent on simply the number of neurons removed from analysis. In contrast, none of these ensembles were involved in trials where shocks appeared to induce a deceleration (**Fig. 2C right, Supp. Table Rows 15-16**).

To address why these ensembles may be differentially necessary for these population dynamics, we compared their activity over the course of shock deliveries between acceleration and deceleration trials (**Fig. 2D**). In the ACC, Shock neurons responded with greater amplitude but desynchronized during shock accelerations (**Fig. 2E, Supp. Table Rows 17,19**). In the RSC, response amplitudes were unchanged between trial types, but Post-shock neurons desynchronized less during shock accelerations (**Fig. 2E, Supp. Table Rows 18, 20**). These results suggest that disorganized but high amplitude shock responses in Shock neurons may drive shock-evoked accelerations in the ACC, while moderate amplitude but more organized activity in Post-shock neurons accelerations in the RSC during unsignaled shock.

### TFC acquisition rapidly induces anticipatory dynamics in RSC neurons

To investigate how these single cell and population dynamics are modified during cued, or expected, foot shocks, 10 ACC- and 10 RSC-implanted mice underwent a five day trace fear conditioning and extinction task as previously described in Rogers et al., 2025 (**Fig. 3A,B, S3A**). Over 8 presentations of foot shock following a 25 sec tone and 20 sec trace period during Acquisition (Acq), mice acquired trace-period freezing as they anticipated the shock, and gradually extinguished this freezing over three days of extinction training (Ext1-3) where they were exposed to six tone presentations (**Fig. 3C, Supp. Table Row 21**).

**Figure 3.**
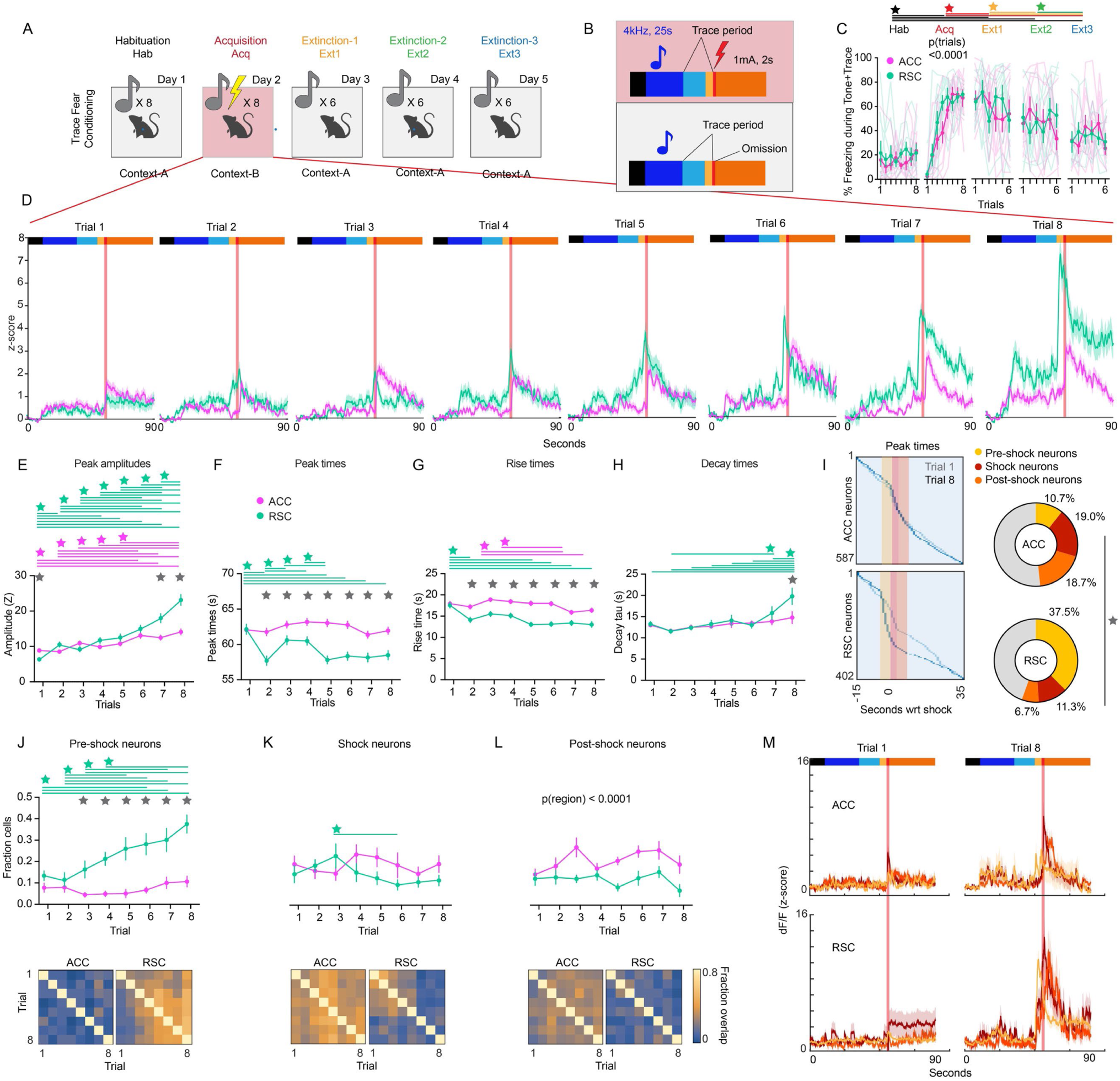
RSC, but not ACC, neurons gradually anticipate shocks during TFC. **A.** Experimental diagram. **B.** Stimulus diagram. **C.** Mean and SEM % freezing during trace period across all sessions and trials (n ACC animals = 10, n RSC animals = 10). **D.** Average and SEM PSTHs aligned to shock delivery over trials 1-8 of Acquisition in the ACC (pink, n=587) and RSC (green, n=402). Two-Way RM ANOVA: pSession < 0.0001, F(4,72) = 33.24. Bars indicate trial periods as color coded in panel B. **E.** Across all ACC and RSC neurons, average peak amplitude of signal during each trial of Acquisition (Two-Way ANOVA: pInteraction < 0.0001, F(7,7896) = 11.01). **F.** Across all ACC and RSC neurons, average time of peak signal during each trial of Acquisition (Two-Way ANOVA: pInteraction = 0.0002, F(1,7896) = 4.07. **G.** Across all ACC and RSC neurons, average rise time of peak signal during each trial of Acquisition (Two-Way ANOVA: pInteraction = 0.017, F(7,7896) = 3.30). **H.** Across all ACC and RSC neurons, average decay of peak signal during each trial of Acquisition (Two-Way ANOVA: pInteraction = 0.044, F(7,7896) = 2.07). **I.** Left: time of peak signal for each ACC (top) and RSC (bottom) neuron during trial 1 (faded) and trial 8 (solid) of Acquisition. Right: proportion of neurons peaking within 5s prior to shock (Pre-shock neurons, yellow), during shock (Shock neurons, red), and within 5s post-shock, (Post-shock neurons, orange) (Chi square test, p < 0.0001, Chi square(3) = 23.05). **J.** Fraction of Pre-shock neurons across animals (Two-Way RM ANOVA: pInteraction = 0.0009, F(7,126) = 3.80). Bottom: Fraction overlap of Pre-shock neurons between each session in ACC (left) and RSC (right). **K.** Same as J for Shock neurons. (Two-Way RM ANOVA: pInteraction = 0.019, F(7,126) = 2.5). **L.** Same as J for Post-shock neurons. (Two-Way RM ANOVA: pRegion < 0.0001, F(1,18) = 27.46). **M.** Mean and SEM PSTHs of each population of neurons in the ACC (top) and RSC (bottom) during Trials 1 (left) and 8 (right) of Acquisition. Stars indicate p<0.05. Green stars indicate change in RSC over time; pink stars indicate change in ACC over time; gray stars with lines indicate change across regions over time; gray stars without lines indicate difference between regions. Statistics in Supplementary Table Rows 21-29.

Overall, the ACC and RSC harbored similar proportions of tone, trace, and shock-responsive neurons with notable exceptions (**Fig. S3B,E,H, Supp. Table Rows 97-106, 108-117, 119-128**). During Acquisition, the fraction of trace upregulated neurons increased and downregulated neurons decreased (**Fig. S3E, Supp. Table Row 108-117**). During Extinction 2, the RSC contained more trace- and shock omission-upregulated neurons than the ACC (**Fig. S3B,H, Supp. Table Row 97-106**). Upregulated neurons with respect to each stimulus were more stable in the RSC than the ACC, and trace downregulated RSC neurons were less stable than those in the ACC (**Fig. S3C,D,F,G,I,J, Supp. Table Rows 107, 118, 129**).

Over the 8 trials of Acquisition, striking differences from the unsignaled shock condition and between regions emerged (**Fig. 3D**). Notably, on average, the RSC population signal began peaking just *prior to* the shock delivery in later trials, while ACC activity continued to peak afterwards. Furthermore, although peak amplitudes were lower in the RSC than ACC during the first trial of Acquisition, similarly to the unsignaled shock condition, they gradually increased over trials until finally the RSC began responding more strongly around the shock than the ACC (**Fig. 3E, S3K, Supp. Table Row 22**). Median peak times also fell dramatically after the first trial in the RSC, reflecting the tendency of many neurons to begin firing earlier over learning (**Fig. 3F, S3L, Supp. Table Row 23**). This result is reflected in decreased rise times in RSC neurons (**Fig. 3G, S3M, Supp. Table Row 24**). Finally, by the eighth trial, RSC neural activity took longer to decay (**Fig. 3H, S3N, Supp. Table Row 25**). In contrast, the only plasticity observed in ACC neurons over trials was an increase in peak amplitude (**Fig. 3E**). Thus, RSC neurons undergo unique and robust changes in responsiveness over trials of trace fear acquisition: responses become greater, earlier, faster to rise, and slower to fall.

Overlaying cumulative distributions of peak times from the first trial on those from the eighth trial highlights the remodeling of RSC peak times and lack thereof in ACC (**Fig. 3I**). By the eighth trial, the RSC hosted significantly distinct proportions of Pre-shock (ACC: 10.7%, n = 63; RSC: 35.7%, n = 150), Shock (ACC: 19.0%, n = 112; RSC: 11.3%, n = 45), and Post-shock (ACC: 16.7%, n = 98; RSC: 6.7%, n = 27) neurons (**Fig. 3I, right, Supp. Table Row 26**). Strikingly, the fraction of Pre-shock neurons monotonically increased over the course of Acquisition trials in the RSC, becoming significantly greater than that in the ACC by the third trial (**Fig. 3J, top, Supp. Table Row 27**). While these neurons did not form a stable ensemble over trials in the ACC, they gradually coalesced into a highly stable ensemble in RSC with nearly 80% overlap between trial 8 and all others (**Fig. 3J, bottom**). Fractions of Shock neurons were steady over trials and similar between groups (**Fig. 3K, top, Supp. Table Row 28**). Interestingly, while this ensemble was highly stable between trials in the ACC, the same was only true in the RSC for the first 5 trials after which overlaps between trials dropped (**Fig. 3K, bottom**). Finally, the ACC contained more Post-shock neurons than the RSC, which were moderately stable across trials (**Fig. 3L, Supp. Table Row 29**). Neural activity in these groups was clearly separable in both regions by the eighth trial (**Fig. 3M**). Together, these results show that the ACC engages stable ensembles of shock-responsive neurons during 8 presentations of cued shock, whereas RSC neuron activity is gradually but robustly remodeled across trials to anticipate cued shocks.

### Distance traveled in neural subspaces depends on TFC session

To investigate how population dynamics in the RSC and ACC might reflect shock expectation during fear acquisition and extinction, we repeated the protocol described in Fig. 1L-M for each session in our TFC protocol. PCA of each session in each region displayed drifting dynamics between each trial and jumps away from the average coordinate of trial during shocks and expected shocks (**Fig. 4A**). To determine whether this drift between trials is an intrinsic or learning-modified property of population dynamics in these brain regions, we calculated the distance traveled between adjacent trials of within each session (**Fig. 4B, Supp. Table Rows 30-35**). We found that trial-to-trial drift in PC1 was independent of training session and brain region. In contrast, drift between trials in PC2 was highly session dependent. In the ACC, PC2 drifted significantly less during Acquisition than all other sessions, suggesting containment of ACC dynamics to a specific subspace in the presence of an aversive stimulus. In contrast, trial-to-trial drift in RSC PC2 was significantly greater than other sessions from Acquisition through Extinction 2, suggesting trajectory motion is learning-dependent. Within-trial drift was significantly greater during Acquisition in the ACC in both PCs during Acquisition, but not altered in RSC PCs. When distance traveled was calculated as a function of the number of trials between two measurements in PC space, we found that distance traveled in PC1 increases monotonically in both regions while distance traveled in PC2 is greater in temporally nearer than further trials. Finally, distance traveled between the shock periods of the end of a given session and beginning of the next did not depend on brain region, but was greater for PC2 between Acquisition and Extinction 1 than between Extinction 2 and Extinction 3. Thus, within-session trial-to-trial movement in PC2 is learning dependent in the RSC, movement within and between trials depends only on shock in the ACC, and overnight movement is greater between shock and non-shock sessions in both regions.

**Figure 4.**
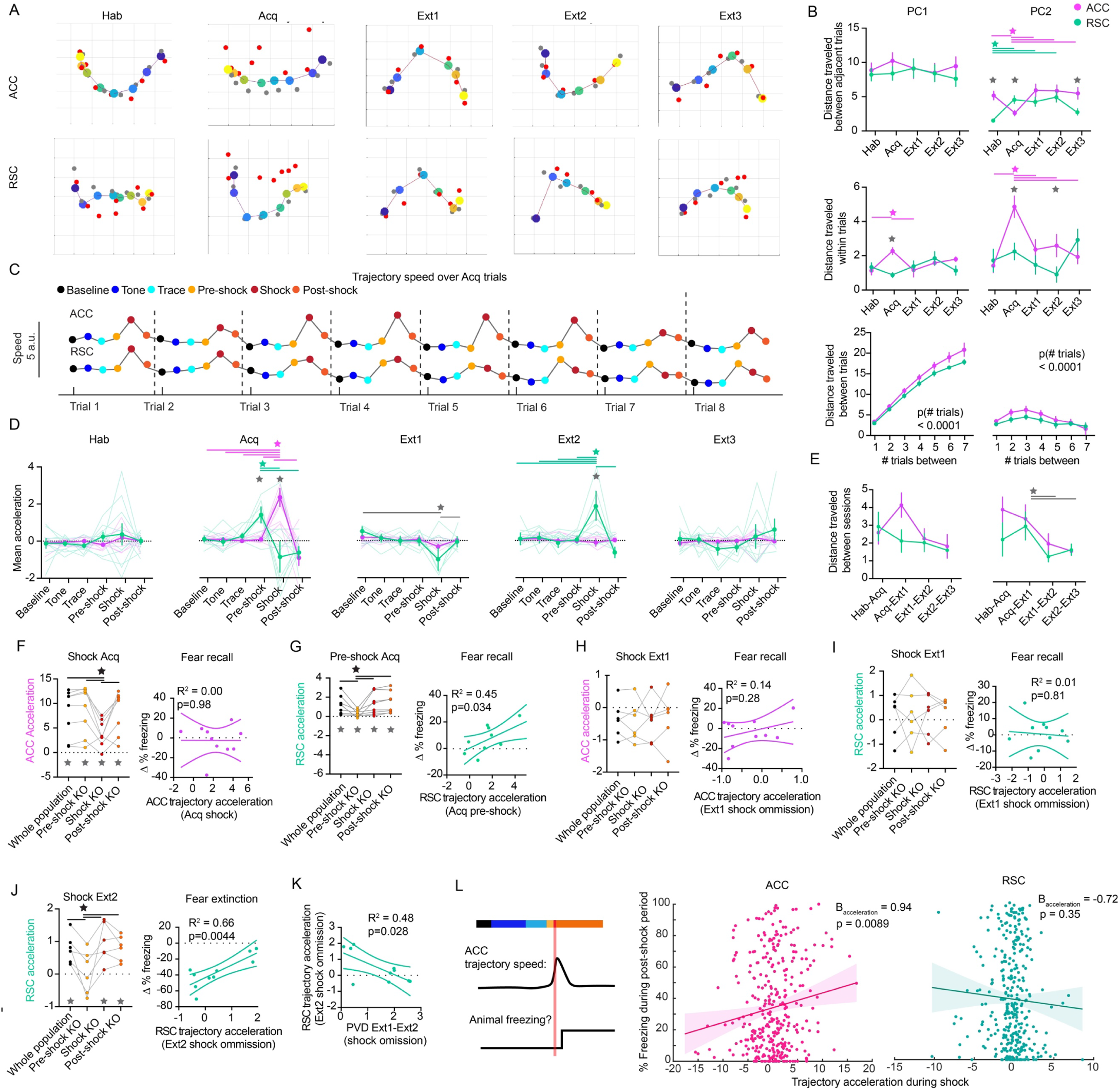
Trajectory accelerations encode decision-making, learning in RSC and ACC, respectively. **A.** PCA of ACC (top) and RSC (bottom) activity over each session (left to right). Large dots are trial averaged coordinates. Dark blue indicates the first trial and bright yellow indicates the last. Gray dots are averaged coordinates during the baseline periods of each trial. Red dots are averaged coordinates during the shocks. **B.** Euclidean distance traveled between (top) and within (bottom) trials of the same session in PC1 (left; Mixed-Effects RM Model: pSession_between_ = 0.044, F(4,138) = 2.52; pInteraction_within_ = 0.0024, F(4,44) = 2.52) and PC2 (right; pInteraction_between_ <0.0001, F(4,138) = 11.23; pInteraction_within_ = 0.012, F(4,44) = 3.55) in ACC (pink) and RSC (green). **C.** Trajectory speed in ACC (top line) and RSC (bottom line) during each trial period throughout Acquisition (baseline = black, tone = blue, trace = cyan, pre-shock = yellow, shock = red, post-shock = orange). **D.** Euclidean distance traveled as a function of the number of trials between measurements in PC1 (left; Mixed-Effects RM Model: pRegion = 0.0002, F(1,188) = 14.10; pNumTrials < 0.0001, F(6, 188) = 202.5) and PC2 (right; Mixed-Effects RM Model: pNumTrials < 0.0001, F(6,188) = 4.38). **E.** ACC and RSC trial-averaged trajectory accelerations during each trial period in each session. Dots and error bars are mean and SEM; lines are individual animals. (Two-Way RM ANOVA: pInteraction_Acq_ < 0.0001, F(5,90) = 8.77; pTrialPeriod_Ext1_= 0.017, F(5,90) = 2.92; pInteraction_Ext2_ = 0.0070, F(5,90) = 3.43). **F.** Euclidean distance traveled between the last and first two shocks/omissions between adjacent sessions in PC1 (left) and PC2 (right; Two-Way RM ANOVA: pSessionPair = 0.0055, F(3,54) = 4.7). **G.** Left: ACC trajectory accelerations during each shock of Acquisition with no, Pre-shock, Shock, or Post-shock neurons removed. Dots are trials, and accelerations are from the pooled PCA (One-Way RM ANOVA: pCondition < 0.0001, F(3,21) = 12.85; One-sample t-tests: pWholePopulation = 0.0017, t(7) = 4.90; pPre-shockKO = 0.0015, t(7) = 5.06; pShockKO = 0.0045, t(7) = 4.11; pPostshockKO = 0.0009, t(7) = 5.49). Right: Linear regression of fear recall (defined as change in % freezing from late Acquisition tone+trace to early Extinction 1 tone+trace) on shock period trajectory acceleration across individual animals. **H.** Same as G for RSC trajectory accelerations during each pre-shock period of Acquisition (One-Way RM ANOVA: pCondition = 0.0008, F(3,21) = 8.28; One-sample t-tests: pWholePopulation = 0.014, t(7) = 3.28; pPre-shockKO = 0.026, t(7) = 2.82; pShockKO = 0.015, t(7) = 3.19; pPostshockKO = 0.0078, t(7) = 3.69). **I.** Same as G for ACC trajectory decelerations during each shock omission of Extinction. **J.** Same as G for RSC trajectory decelerations during each shock omission of Extinction 1. **K.** Left: RSC trajectory accelerations during Fig. 3 **continued.** each shock omission of Extinction 2 with no, Pre-shock, Shock, or Post-shock neurons removed. Dots are trials, and accelerations are computed from the pooled PCA (One-Way RM ANOVA: pCondition = 0.0015, F(3,15) = 8.54; One-sample t-tests: pWholePopulation = 0.0075, t(5) = 4.32; pPre-shockKO = 0.79, t(5) = 0.28; pShockKO = 0.011, t(5) = 3.94; pPostshockKO = 0.0036, t(5) = 5.14). Right: Linear regression of fear extinction (defined as change in % freezing from early Extinction 1 tone+trace to early Extinction 3 tone+trace) on shock period trajectory acceleration across individual animals. **L.** Linear regression of shock period trajectory acceleration on Euclidean distance traveled during the last two shocks of Extinction 1 to the first two shocks of Extinction 2. **M.** Linear mixed model predicting freezing during the post-shock period from the trajectory acceleration in the shock period immediately prior (diagrammed on the left) in the ACC (middle) and RSC (right), with acceleration treated as a fixed effect and animals as random effects. Dots are each a given trial in a given animal. Stars indicate p<0.05. Green stars indicate change in RSC over time; pink stars indicate change in ACC over time; gray stars with lines indicate change across regions over time; gray stars without lines indicate difference between regions. Statistics in Supplementary Table Rows 30-75.

### RSC trajectory accelerations precede expected shocks or omissions over TFC

Over trials of Acquisition, trajectory speed followed patterns recapitulating our observations at the single cell level (**Fig. 4C**). While ACC trajectory speed peaked during each shock, peaks in RSC speed gradually shifted earlier such that peak speed was achieved during the pre-shock period. Consistently, during Acquisition, we found, on average, that RSC trajectories accelerated during the pre-shock period, while ACC trajectories accelerated during the shock (**Fig. 4D, Supp. Table Rows 36-40**). Accordingly, distance traveled between shock or omission periods between each session was greatest between Acquisition and Extinction 1 in PC2 than all other pairs (**Fig. 4E, Supp. Table Rows 41-42**). Still, the predictability of the shock by temporally related cue restructures single cell and population dynamics in the RSC and not ACC.

If the expectation of shock is sufficient to drive trajectory accelerations in the RSC, we hypothesized that accelerations around the shock omission period should be observed during early extinction. In contrast, during the shock period of Extinction 1, both regions displayed shock period *decelerations*. Instead, trajectory accelerations in RSC were apparently recovered during the shock omission period of Extinction 2, before vanishing again at Extinction 3. Thus, RSC trajectory accelerations occur when mice are learning to predict a shock at a certain time and when mice are learning to expect its omission, but not when mice are recalling an expectation.

### Pre-shock neurons drive learning-predictive trajectory accelerations in RSC

We then proceeded to systematically test which neural ensembles are involved each trajectory acceleration in each region, and whether these trajectory accelerations predicted learning as measured by change in percent freezing between sessions. In the ACC, Shock neurons were required for Acquisition shock period accelerations, but these accelerations did not predict fear recall (average change in freezing during tone and trace between the last two trials of Acquisition and the first two trials of Extinction 1). Thus, while ACC dynamics encode an aversive stimulus, they do not appear to be related to fear learning (**Fig. 3F, Supp. Table Rows 43-48**). In contrast, in the RSC, Pre-shock neurons were necessary for Acquisition pre-shock accelerations, and these accelerations predicted fear recall (**Fig. 4G, Supp. Table Rows 49-54**). Thus, the strength of shock-predicting trajectories in the RSC predicts how well an animal learns fear, and this Pre-shock relies on a subgroup of neurons that peak before the shock.

In neither brain region trajectory decelerations during the shock omission period of Extinction 1 predictive of fear recall nor reliant on any previously defined group of neurons (**Fig. 4H,I, Supp. Table Rows 55-66**). Finally, we investigated the trajectory acceleration in the RSC during the shock omission of Extinction 2 (**Fig. 4J, Supp. Table Rows 67-72**).

Surprisingly, these accelerations were in fact dependent on Pre-shock neurons, and, furthermore, predicted fear extinction (defined as the average change in % freezing between the tone and trace periods of the first two trials of Extinction 1 and the first two trials of Extinction 3).

Importantly, this relationship was positive: greater shock omission accelerations predicted a less negative change in freezing over extinction, or less learning. If shock omission trajectory accelerations indicate the acquisition of an expectation about shock omission, one would expect this association to be reversed. Importantly, though, all mice did *reduce* their freezing between these two timepoints, so these accelerations must not represent reinforcement of the fear memory. We therefore wondered if these accelerations still represented prediction of shock omission but that, nevertheless, this was a less optimal strategy of extinguishing fear.

If this hypothesis is correct, then mice that underwent lesser trajectory accelerations during the shock omission period of Extinction 2 should have undergone greater remodeling of shock period representations between Extinction 1 and Extinction 2 – that is, overnight. Indeed, the greater the population vector distance between the last two shock omissions of Extinction 1 and the first two of Extinction 2, the greater were animals’ trajectory accelerations during Extinction 2 shock omission (**Fig. 4K, Supp. Table Row 73**). This evidence circumstantially supports the hypothesis that animals employ distinct, though not mutually exclusive, strategies to extinguish fear, with distinct levels of efficacy: online learning, whereby the expectation of shock omissions are actively represented mid-extinction in the RSC, and offline learning, whereby the absence of shocks during Extinction 1 is consolidated overnight.

### Trajectory accelerations predict decision to freeze in ACC

In Figs. 3 and 4, we have shown that the ACC appears to have specialized single cell and population dynamics for shock presentation independent of expectation. Although we also showed that the ACC does not appear to display dynamics important for fear learning or extinction, the ACC has been previously shown to be crucial for decision-making and behavioral-selection. We therefore wondered whether trajectory accelerations in the ACC, whether they occur during a shock or shock omission, would be associated a subsequent decision to freeze or not to freeze. To test this, we built a linear mixed model, in which biological replicates were treated as random fixed effects, to predict freezing during the post-shock period from trajectory accelerations during the shock period in individual trials from all sessions (**Fig. 4L, Supp. Table Rows 74-75**). We found that shock/omission trajectory accelerations significantly predicted subsequent freezing in the ACC, but not the RSC. Thus, while RSC dynamics are predictive of fear learning and extinction over minutes and days, trajectory accelerations in the ACC are predictive of the choice to freeze on the order of seconds.

### Changes in neural synchrony and peak time may explain contribution to trajectory accelerations

We then explored what rendered particular ensembles capable of inducing trajectory accelerations during the shock/omission period in each region (**Fig. 5A**). In the ACC, peak times for all ensembles were earlier during Pre-shock than all other sessions, but ensembles were not distinguished (**Fig. 5B, Supp. Table Rows 76-77**). In the RSC, the same held, except Pre-shock neurons peaked significantly later during Extinction 2 than the other ensembles. In both regions, peak amplitudes were greater on average during Acquisition than other sessions, but ensembles were not distinguished (**Fig. 5C, Supp. Table Rows 78-79**). Interestingly, though, ensembles differed substantially as to how much more or less synchronous they became during the shock period compared to baseline (**Fig. 5D, Supp. Table Rows 80-81**). In the ACC, Shock neurons significantly desynchronized more than other ensembles during Acquisition shock. In the RSC, Pre-shock neurons desynchronized less than both others during Acquisition shock and became more synchronous than both others during Extinction 2. Therefore, while the Acquisition shock period acceleration in the ACC appears related to the desynchronized but high amplitude and early peaking of Shock neurons, the Extinction 2 omission period acceleration in the RSC appears to be related to synchronized activity preceding a late peak in Pre-shock neurons.

**Figure 5.**
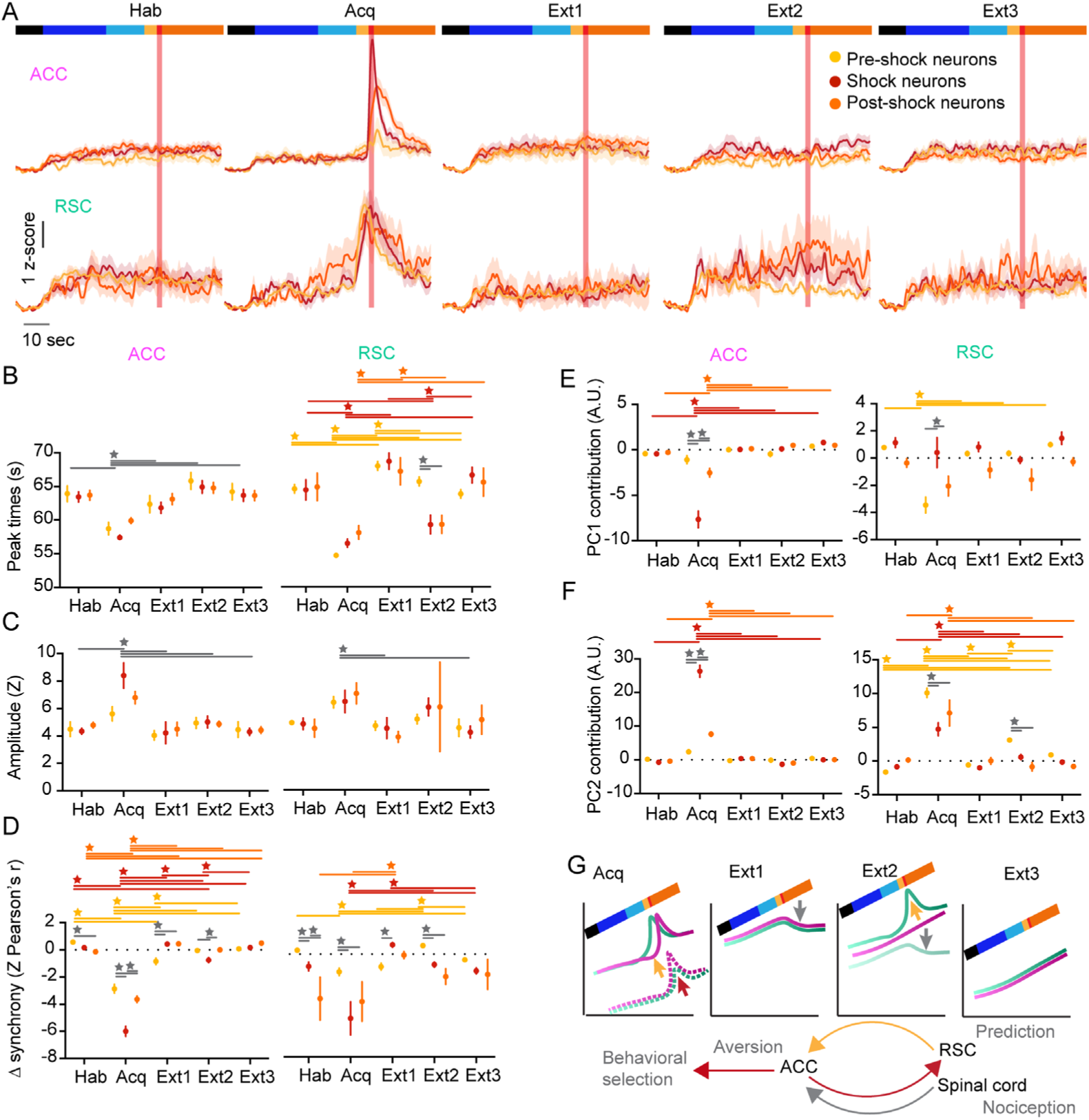
Properties of ensembles explain their contributions to population dynamics. **A.** Mean and SEM PSTHs of each population of neurons in the ACC (top) and RSC (bottom) during over each session. **B.** Across each neural population, average time of peak signal during each session in ACC (left; Two-Way ANOVA: pSession < 0.0001, F(8,1340) = 0.60) and RSC (right; Two-Way ANOVA: pInteraction < 0.0001, F(8,1170) = 5.11). **C.** Across each neural population, average amplitude of peak signal during each session in ACC (left; Two-Way ANOVA: pSession < 0.0001, F(4,1340) = 14,71) and RSC (right; Two-Way ANOVA: pSession = 0.0065, F(4,1170) = 3.60). **D.** Across each neural population, average change in synchrony (z-score Pearson’s r) in ACC (left; Two-Way ANOVA: pSession < 0.0001, F(4,1340) = 339.10) and RSC (right; Two-Way ANOVA: pInteraction < 0.0001, F(8,1170) = 8.71). **E.** Across each neural population, average contribution to PC1 during shock (contribution of neuron k in PC1 during shock = *week_,1_\**A_k,shock_) in ACC (left; Two-Way ANOVA: pInteraction < 0.0001, F(8,1340) = 19.10) and RSC (right; Two-Way ANOVA: pInteraction = 0.0002, F(8,1179) 3.89). **F.** Across each neural population, average contribution to PC2 during shock (contribution of neuron k in PC1 during shock = *week_,1_\**A_k,shock_) in ACC (left; Two-Way ANOVA: pInteraction < 0.0001, F(8,1340) = 94.43) and RSC (right; Two-Way ANOVA: pInteraction < 0.0001, F(8,1170) = 15.20). **G.** Diagram of results. During Acquisition, Pre-shock neurons drive pre-shock acceleration in the RSC, and Shock neurons drive shock-acceleration in the ACC. During Extinction 1, an unknown population of neurons drive omitted-shock decelerations in the RSC and ACC. During Extinction 2, Pre-shock neurons drive shock accelerations in the RSC of poorly extinguishing mice, but some other ensemble drives decelerations in the RSC of well extinguishing mice. During Extinction 3, there are no more trajectory accelerations. The precise timing of these events, that evolves over trials in Acquisition, suggests a feedback loop: shock neurons in the ACC could be relaying aversive information to the RSC after initial shocks, which then uses the cue to begin predicting the shock and influence freezing and behavioral selection in future sessions via the ACC. Stars indicate p<0.05. Green stars indicate change in RSC over time; pink stars indicate change in ACC over time; gray stars with lines indicate change across regions over time; gray stars without lines indicate difference between regions. Statistics in Supplementary Table Rows 76-85.

We further supposed that the neural ensembles that drive trajectory accelerations during shocks/omissions should explain more of the PCs in that session than other ensembles. Contribution to a given PC was calculated by multiplying the weight of a given neuron in that PC by the sum of its activity during the shock. Indeed, in the ACC, Shock neurons most strongly reduced PC1 and increased PC2 during the shock, but not other sessions (**Fig. 5E,F, Supp. Table Rows 82-85**). In the RSC, Pre-shock neurons reduced PC1 the most during Acquisition shock, but not more than Post-shock neurons (**Fig. 5E**). These neurons also most strongly contributed to PC2 during both Acquisition shock and Extinction 2 omission.

In sum, the RSC undergoes specialized dynamics during trace fear conditioning and extinction, distinct from repeated, unsignaled foot shock exposure and from the ACC (**Fig. 5G**). While both regions initially respond to foot shock following the first cue presentation, over fear conditioning, RSC trajectory accelerations begin to precede shock, a shift driven by the gradual recruitment of Pre-shock neurons. Overnight, a large population vector distance is traversed in PC space in both regions, a displacement to a new subspace that perhaps explains why shock omissions in Extinction 1, during fear recall, evoke decelerations. In the RSC, in mice that will go on to extinguish well, large population vector distance is traversed as well. In mice that go on to extinguish poorly, shock omission period trajectory accelerations occur in Extinction 2. Finally, omission accelerations vanish during Extinction 3. Together, the temporal structure of this evidence suggests a potential circuit that could govern fear learning and recall: the ACC receives ascending nociceptive input that Shock neurons relay to the RSC to evoke its initial small responses; then the RSC learns the cue and comes to predict the shock via its Pre-shock neurons; finally, this Pre-shock signal is then relayed back to the ACC to evoke freezing, whether in the presence or absence of a genuine shock.

## Discussion

By leveraging longitudinal calcium imaging of the ACC and RSC across both unsignaled shock and trace fear conditioning paradigms, we identify how distinct specializations of these inter-connected regions shape aversive behaviors and learning. In both regions, we observed stable ensembles and population-level dynamics in response to unsignaled shocks across hours to days, supporting the presence of persistent nociceptive representations. However, only the RSC exhibited robust modulation of its activity when shocks were preceded by a cue, gradually recruiting an ensemble peaking 0-5sec prior to the shock. This distinction suggests a division of labor within the cingulate continuum in which the ACC encodes ongoing nociceptive information with relative stability to guide immediate behavioral decision-making, while the RSC flexibly reshapes its dynamics to capture the temporal structure of harmful events and thereby guide learning.

The stability of shock-responsive ensembles in both ACC and RSC may help resolve a long-standing debate about whether the ACC harbors a persistent nociceptive population. Both regions maintained populations of Shock neurons stable across hours and, to a lesser degree, across days, consistent with the notion that ascending nociceptive signals are durably represented in cortical circuits. Importantly, however, ensemble composition diverged over time. The ACC Post-shock neurons displayed high stability, while RSC Post-shock neurons only coalesced into stable populations after a week. This delayed recruitment suggests that RSC does not simply encode aversive experiences in real time, but gradually integrates the experience into representations that persist over longer timescales, echoing its role in remote and contextual memory.

At the population level, neural trajectories in both regions exhibited accelerations at shock onset, but their ensemble underpinnings differed. In the ACC, accelerations were driven by Shock neurons whose responses were high in amplitude but disorganized in synchrony. In the RSC, by contrast, accelerations depended on Post-shock neurons whose responses were moderate in amplitude but more temporally organized. These results suggest that shared dynamics at the level of neural state space, such as trajectory accelerations, can be implemented by distinct local circuits across regions, reflecting their differing computational specializations.

During TFC, the divergence between regions became most apparent. ACC trajectories consistently accelerated at the shock itself, and these accelerations predicted the immediate decision to freeze on a trial-by-trial basis across sessions. RSC trajectories, in contrast, shifted progressively earlier across Acquisition trials, ultimately accelerating during the pre-shock period. This remodeling was driven by the gradual recruitment of stable ensembles of Pre-shock neurons. The strength of these anticipatory dynamics predicted how well mice acquired fear, highlighting the RSC as a cortical locus for linking temporally separated events. These findings resonate with prior work implicating the ACC in action selection and immediate aversive processing, and the RSC in spatiotemporal integration and episodic memory.

The extinction data further underscore the RSC’s role in encoding predictions about aversive events and their absence. During early extinction, both regions displayed trajectory decelerations at the time of expected shock. By Extinction 2, RSC anticipatory ensembles again drove trajectory accelerations, this time at the omission of shock. Intriguingly, these omission-related accelerations predicted poorer extinction and less change in the shock representation overnight. These results point to multiple, possibly competing, neural strategies for fear suppression: offline restructuring of representations overnight versus online encoding of omissions within session. If fear learning corresponds to accelerations, fear recall to decelerations, and extinction learning to accelerations or large representational drift, then these shock-period traversals over neural subspace likely reflect contingency learning rather than aversion itself, a large functional divergence from the unsignaled shock paradigm. Perhaps the lack of contingency in the unsignaled shock experiment explains the dissolution of trajectory accelerations at 2weeks, due to contextual fear associations^41^.

Mechanistically, ensemble-level synchrony emerged as a key indicator of which ensembles might be crucial to trajectory accelerations. In the ACC, desynchronization of Shock neurons was occurred during accelerations, likely reflecting divergent roles broadcasting nociception or fear and guiding behavioral strategies. In the RSC, Pre-shock neurons synchronized during omission accelerations, pointing to a circuit mechanism by which predictions about temporally structured stimuli are encoded.

The non-overlapping connectivity between these regions frames these disparate operations in aversive learning. The ACC is reciprocally connected to the amygdala, periacqueductal gray, and ventral striatum^47–50^. The RSC is reciprocally connected to hippocampus, entorhinal cortex, and subiculum^51–54^. Therefore, while the ACC is anatomically optimized to integrate and transform information about valence, salience, motivation, and behavioral strategy, the RSC is anatomically optimized to integrate and transform information about memory, time, space, and context. In the future, it would be of interest to perform circuit interrogation studies to explore how pre-shock predictions in the RSC might be routed to the ACC to induce an aversive state, which could then instruct the periacqueductal gray to execute freezing or escape strategies as previously described^55,56^.

Together, these findings identify a cortical network orchestrating how aversion is experienced, learned about, and acted upon. The ACC stably encodes nociceptive input and shapes immediate behavioral selection, while the RSC transforms these signals into temporally structured representations that guide the acquisition and extinction of fear. This division of labor reconciles the otherwise disparate functional specializations of the two regions and positions their interaction as a key substrate for transforming aversive sensation into adaptive learning and memory.

## Acknowledgments

This work was supported by: National Institute of Health (NIH) awards NIGMS DP2GM140923 (G.C.) and NINDS F31NS143421 (S.A.R.). We thank Jacqueline Wu and the Penn University Laboratory Animal Resources (ULAR) veterinarians and husbandry staff for taking care of animals used in this study.

## Code, data, materials availability

Code and pre-processed data will be available at https://github.com/sarogers9/ACCRSC_rogersEtAl2025. Raw data files are available upon request.

## Author contributions

Conceptualization: S.A.R, G.C.

Data Collection: S.A.R., C.S.O., L.L.E.

Data Analysis: S.A.R.

Software: S.A.R.

Writing and editing: S.A.R., G.C.

Funding: S.A.R., G.C.

## Methods

### Experimental methods

#### Ethical statement

All research complies with all relevant ethical regulations, including those established by the University of Pennsylvania Institutional Animal Care and Use Committee and Institutional Biosafety Committee.

#### Animals

Animals used in all studies were C57BL/6J mice from Jackson Laboratories (research resource identifier— IMSR_JAX:000664). Mice were kept on a reverse 12hr light/12hr dark cycle. Behavior was performed at least 1hr and no more than 5hours following lights-off. Males of 8-10 weeks of age underwent viral injection surgeries, followed by implantation of 4.0 (length) × 1.0 mm (diameter) GRIN lens at 10–12 weeks, and behavior at 12– 16 weeks (minimum 2-week recovery time from last surgery). Mice were singly housed following implant surgery. The ambient temperature in the rodent colony room was ∼70 °F, and the room humidity was ∼46%.

#### Sample size

For both experiments, sample sizes were set to have >100 longitudinally registered neurons per region to appropriately explore population dynamics. For TFC, we aimed to have at least ten animals per region to relate neural activity outcomes to behavior. In the unsignaled shock experiment, there were three RSC- and four ACC-implanted animals (147 and 596 longitudinally registered neurons, respectively). In the TFC experiment, there were ten animals for each region (402 and 587 longitudinally registered neurons, respectively).

#### Randomization

Animals were randomly assigned to treatment groups before the experiment.

#### Blinding

Investigator was informally blinded during the experiment, such that data were not analyzed until data collection was complete and extraction of behavior and neural activity traces was complete. Investigator checked treatment conditions once it became necessary to assign animal data to treatment groups.

#### Unsignaled shock

One week before behavioral testing, GRIN lens-implanted mice were habituated to the Miniscope for 2 days in 10-min sessions in the home cage. All mice underwent behavioral training and testing in Med Associates fear conditioning boxes over five sessions, at 0hours, 4hours, 24hours, 1week, and 2week timepoints. Med Associates fear conditioning chambers (ENV-307W) contained grid floors, under which lay a tray with paper towels. Each session, mice were exposed to eight, two second, 1 mA shocks 105±10 seconds apart. Intertrial intervals were pseudorandomly jittered.

#### TFC conditioning and extinction

One week before behavioral testing, GRIN lens-implanted mice were habituated to the Miniscope for 2 days in 10-min sessions in the home cage. All mice underwent behavioral training and testing in Med Associates fear conditioning boxes for 5 days. In context A, Med Associates chambers were equipped with smooth white floor inserts and cleaned with ethanol to provide a unique olfactory, tactile and visual context. In context B, the shock grid floor was exposed, paper towels were placed in a tray under the floor, and chambers were cleaned with Clidox. The 5 days of behavioral testing consisted of Habituation (Hab), Acq and Ext1–Ext3. Hab and Ext1– Ext3 took place in context A, and Acquisition took place in context B. The CS consisted of a 4 kHz, 75 dB tone delivered in 25, 200-ms pips at 1 Hz. During Acquisition, the CS was followed by a 20-s trace period preceding a 1 mA, 2-s shock. On all other days, the shock was omitted. Habituation and Acquisition consisted of eight trials, with jittered intertrial intervals (ITIs) of 60 ± 10 s. During Extinctions 1–3, there were six trials per session.

#### Miniscope recordings

Before each session, the Inscopix miniscope was magnetically attached to the mouse’s implanted baseplate and screwed into place. During the sessions, recordings were remotely controlled and streamed to a laptop for live monitoring. Recordings were made at LED power (0.7–1.5 mW), gain (1.0–3.0) and focus (0–300 μm) settings deemed appropriate for each mouse and kept as consistent between recording sessions as possible.

#### Viral injections

For Miniscope studies, all mice were unilaterally injected with 800 nl of AAV9-syn-GCaMP8m-WPRE at a titer of 1.2 × 10−12 (Addgene virus, 162375) in the ACC or RSC. Coordinates were chosen from past studies (ACC: anterior-posterior (AP)= +1.50, medial-lateral (ML) = +0.3 and dorsal-ventral (DV) = −1.0; RSC: AP = −2.25, ML = +0.3 and DV = −0.8). Mice were anesthetized with isoflurane. Hair was removed with Nair, and the skin was sterilized with Betadine and ethanol. An incision was made with scissors along the scalp. Tissue was cleared from the skull surface using an air blast. The skull was leveled such that the bregma–lambda and ML DV difference was within ±0.1 mm. A craniotomy was made at the chosen coordinates with a dental drill. A needle was lowered to the target coordinates through the craniotomy, and the virus was infused at 100 nL/min. The needle was left in the brain 10 min after infusion before being slowly withdrawn. The incision was sutured, and the animal was administered meloxicam before being placed under a heat lamp for recovery.

#### Miniscope implantation

These surgeries subsequently followed the same protocol until the craniotomy step. A 1 mm craniotomy was made by slowly widening the craniotomy with the dental drill. Dura was peeled back using microscissors, sharp forceps and curved forceps. The craniotomy was regularly flushed with saline, and gel foam was applied to absorb blood. An Inscopix Pro-View Integrated GRIN lens and baseplate system was attached to the Miniscope and a stereotax. Using the Inscopix stereotax attachment, the lens was slowly lowered into a position over the injection site. The final DV coordinate was determined by assessing the view through the Miniscope stream. If tissue architecture could be observed in full focus, the lens was implanted at that coordinate (ACC: -1.2 to -1.8 DV; RSC: -0.6 to -0.3 DV). The GRIN lens + baseplate system was secured to the skull with Metabond and then dental cement. After surgery, mice were singly housed and injected with meloxicam for three consecutive days during recovery.

#### Miniscope validation

Before admission to the experiment, the miniscope was magnetically attached to each animal’s implant for habituation and streamed using Inscopix Data Acquisition Software. If many cells could be observed during spontaneous behavior in the home cage, the mouse was admitted. If only a few cells were visible, the session was recorded and analyzed in Inscopix Data Processing Software (IDPS) to determine the number of observable cells. If an animal had >20 identifiable cells, they were admitted into the study. Others were sacrificed.

#### Exclusions

No animals excluded from the study.

### Analytical methods

#### Behavioral

For the TFC experiment, behavior was recorded by Basler cameras into Pylon Viewer at 15 Hz. Videos were then processed in the open-source ezTrack Jupyter Notebook^57^. The algorithm was calibrated to the standard light fluctuations in the empty chambers and the empty chambers with the Miniscope wire dangling in them for each respective study. A freezing threshold was determined in terms of the number of pixels changed per frame by visually validating portions of videos classified as ‘freezing’ or ‘ moving’ by the algorithm. A freezing threshold of 150 pixels per frame was used. An animal was only classified as ‘freezing’ if the pixels per frame remained below the threshold for at least 1 sec, or 15 frames. Freezing status per frame was exported in a CSV file and postprocessed in MATLAB to calculate the percentage freezing windows of time. Freezing videos were aligned to trial times by beginning analysis at the first frame of the red light in the Med Associates boxes switching on, indicating session start.

#### Calcium imaging preprocessing

Videos were downloaded from the Inscopix Data Acquisition Box and uploaded to IDPS. Videos were spatially downsampled by a factor of 4 and spatially band-pass filtered between 0.005 and 0.500. Videos were then motion corrected with respect to their mean frame. Cells were identified and extracted using CNFM-E (default parameters in the Inscopix implementation of CNMF-E, except the minimum in-line pixel correlation = 0.7 and minimum signal-to-noise ratio = 7.0) and second-order deconvolved using SCS. Videos across 5 days of behavioral training were longitudinally registered in IDPS (minimum normalized cross-correlation = 0.1). Normalized cross-correlation score and centroid distance from Habituation were used to validate registration. Only cells registered on all 5 days were considered for further analysis.

#### Calcium imaging postprocessing

Most subsequent analysis was performed in custom MATLAB scripts (version 2020a), available in the associated GitHub. Deconvolved calcium traces of cells from each session were aligned according to their global cell index determined in longitudinal registration. As the window considered for each trial included a 10 sec baseline period, a 25 sec stim period, a 20 sec trace period, a 2 sec shock/omission period and 30 sec after, each trial was 90 sec. Miniscope recordings were started exactly 30.00 s before behavioral session start, and this information was used to align data to behavior and neural data.

#### Ensemble characterization

For all neurons, their peak activity during a trial was identified as the maximum of the z-score dF/F value after from 15 sec before to 30 sec after the shock or omitted shock. Peak amplitude and time on each trial were calculated with the findpeaks function in Matlab. Rise time was defined as the time from the previous local minimum activity to the peak itself. Decay times were calculated by finding the frame at which a neuron’s activity was ≥ Activity_baseline_ + (Activity_peak_ - Activity_baseline_)*e^-1/2^. Pre-shock, Shock, and Post-shock neurons were defined as those neurons peaking within the 5 sec before, during, and 5 sec after the shock, respectively, on the last trial of 0hours in the unsignaled shock experiment or of Acquisition in the TFC experiment. Stability of these ensembles was assessed by assessing the overlap of the neurons peaking at these times between trials in sessions containing shocks.

#### Trajectory accelerations

To calculate trajectory accelerations, PCA was performed on the z-scored activity of all longitudinally registered neurons, concatenated over trials, from an individual animal within a particular session. Trajectory speed at each trial periods was calculated as the first derivative of the position in the first two PCs during the first 5 seconds of each trial period, or the two second shock (baseline: 0-4 sec, tone: 10-14 sec, trace: 36-40 sec, pre-shock: 50-54 sec, shock: 55-58 sec, and post-shock: 59-63 sec). Acceleration was calculated by taking the second derivative.

To test the effect of removing neural ensembles, the same analysis was done after pooling neurons from all animals implanted in a given region and statistics were done using trials, rather than animals, as replicates. This procedure circumvented the issues caused by removing different proportions of neurons from each animals. For instance, removing 60% of neurons from one animal and 20% of neurons from another would differently affect trajectory acceleration computation (namely, removing 60% of neurons would compress PC space more than removing 20%) and confound results. The protocol was repeated while systematically removing each ensemble.

#### Distance traveled metrics

To calculate the distance traveled in PC space between trials, the Euclidean distance was taken from the average coordinate of a given trial to that of its adjacent trials. To calculate the same quantity within trials, the Euclidean distance was taken from the average coordinate of the first half of a given trial to that of latter half. To calculate distance traveled between shock and shock omission representations across days, the Euclidean distance was taken from the average coordinate of the last two shocks or omissions of a given day to the first two of the next day.

#### Correlating neural activity with behavior

To determine the relationship between trajectory accelerations and learning, accelerations during the pre-shock, shock, or shock omissions were smoothed by 20 frames (1 sec) and averaged over the session of interest for each animal. Fear recall was defined as the change in average percent freezing from the tone and trace of the last two trials of Acquisition to the that of the first two trials of Extinction 1, such that near zero or positive values indicate greater recall and very negative values indicate incomplete recall. Fear extinction was defined as the change in average percent freezing from the tone and trace of the first two trials of Extinction 1 – where fear recall can be assessed – and the first two trials of Extinction 3 – where extinction recall can be assessed. Fear recall was linearly regressed on trajectory accelerations from the pre-shock period of Acquisition (RSC), the shock period of Acquisition (ACC), and the shock omission period of Extinction 1 (both) in Prism. Fear extinction was linearly regressed on trajectory accelerations from the shock omission period of Extinction 2 (RSC).

To determine the relationship between trajectory accelerations and the decision to freeze, shock period accelerations were calculated for each trial and each animal in a given region, and the percent freezing was calculated during the post-shock period for each. To account for the repeated observations from biological replicates, we built a linear mixed model where percent freezing was modeled as the response variable, accelerations as the fixed effect, and animal label as a random effect.

#### Change in neural synchrony

For each ensemble, raw neural activity was smoothed over 20 frames (1 sec) and the lower half of their correlation matrix from 20 frame sliding windows was averaged. To calculate the change from baseline, correlation values were z-scored to baseline.

#### PC contribution

Contribution of each neural ensemble to each PC during the shock was calculated by multiplying a given neuron’s coefficient withing that PC by its average z-scored activity during the shock. This value was then averaged over neurons.

#### Statistics

Statistics were calculated in Prism. All between-region or between-ensemble comparisons over time were Two-Way RM ANOVAs or Mixed Effect Models (when there was missing data, e.g. NaNs). When removing neural ensembles from trajectory accelerations, One-Way RM ANOVAs were used to test effect of condition and one-sample t-tests were used to test if accelerations were different from zero.

#### Exclusions

For the linear mixed model, 5 ACC shock accelerations were 1-2 orders of magnitude higher than all others, and the iterative Grubb’s test indicated these were outliers. These observations were excluded from the model.

## Supplementary figures

**Fig. S1.**
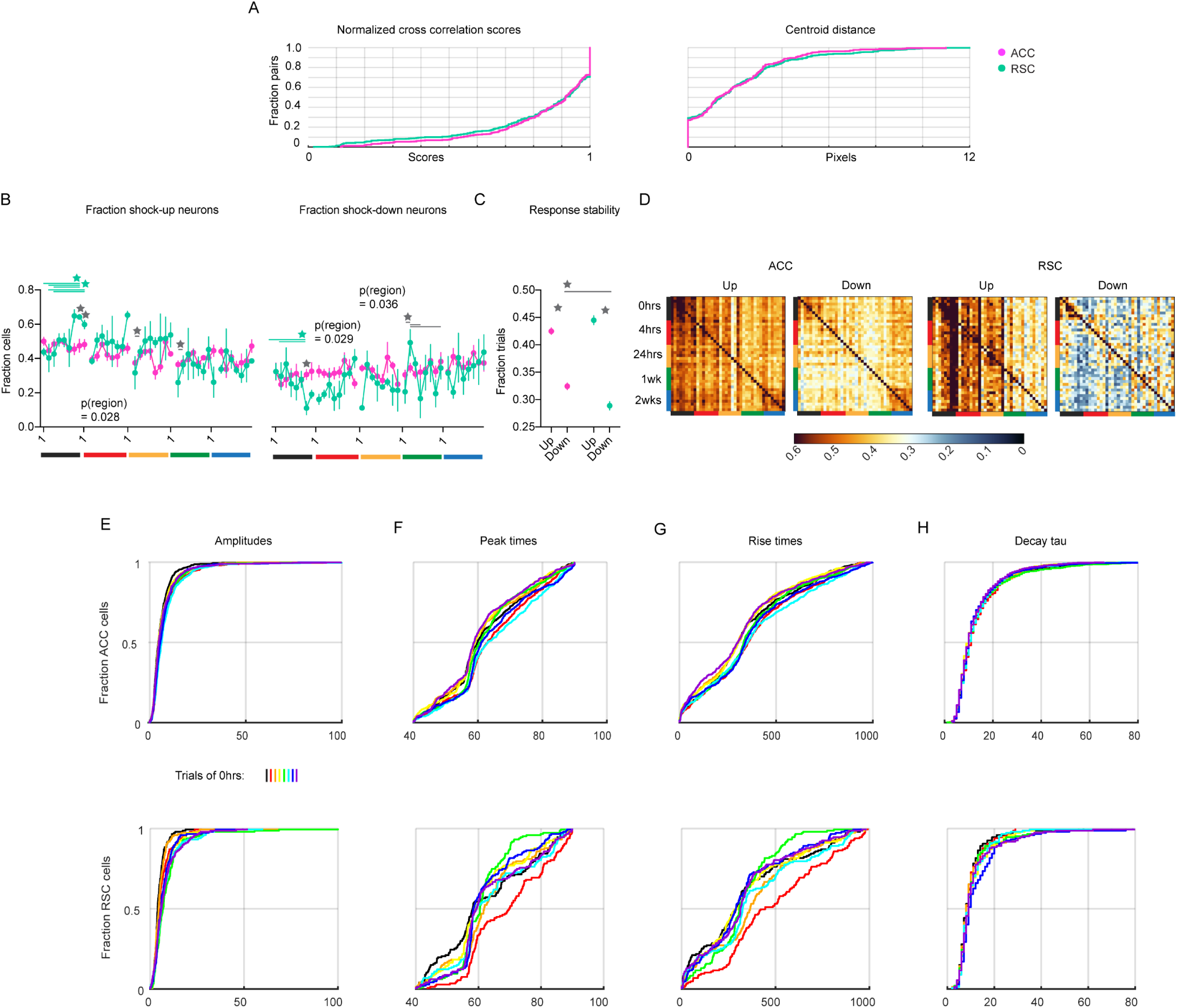
Neural tuning over unsignaled shock. **A.** Cumulative distributions of normalized cross-correlation scores (left) and centroid distances (right) between longitudinally registered ROIs in ACC (pink) and RSC (green). **B.** Fraction of significantly shock-upregulated (left; Two-Way RM ANOVA: pInteraction_0hours_ = 0.023, F(7,35) = 2.73; pRegion_4hours_ = 0.037, F(1,5) = 7.95; pTrial_1week_ = 0.028, F(7,35) = 2.61) and -downregulated (right; Two-Way RM ANOVA: pInteraction_0hours_ = 0.025, F(7,35) = 2.67; pRegion_4hours_ = 0.029, F(1,5) = 9.27; pRegion_24hours_ = 0.036, F(1,5) = 8.14; pTrial_1week_ = 0.0050, F(1,5) = 3.61) neurons in each trial within each session. Colored bars indicate session ID. **C.** Fraction of trials that significantly modulated neurons maintained their response properties (Two-Way ANOVA; pInteraction = 0.0020, F(1,1482) = 9.55). **D.** Heatmaps showing the overlap of significantly up- and down-regulated neurons in the ACC (left) and RSC (right) over trials. Colored bars indicate session ID. **E.** Cumulative distributions of response amplitudes of ACC (top) and RSC (bottom) neurons over each trials, colored according to black-ROYGBIV from trials 1-8 of the first session. **F.** Same as E for peak times. **G.** Same as E for rise times. **H.** Same as E for decay times. Stars indicate p<0.05. Green stars indicate change in RSC over time; pink stars indicate change in ACC over time; gray stars with lines indicate change across regions over time; gray stars without lines indicate difference between regions. Statistics in Supplementary Table Rows 86-96.

**Fig. S2.**
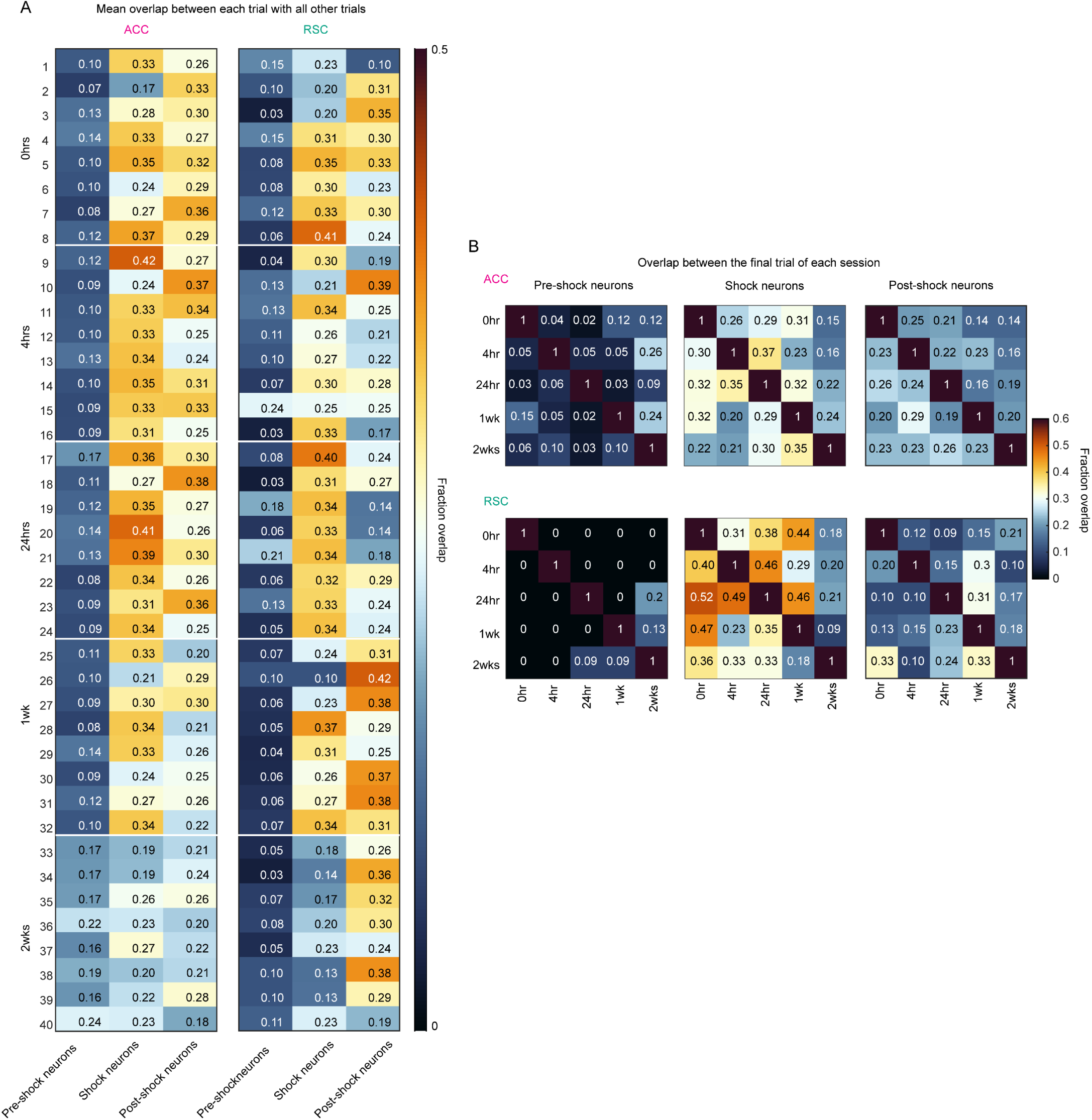
Ensemble overlap quantification. **A.** Mean overlap of Pre-shock, Shock, and Post-shock ensembles between each trial with all other trials in ACC (left) and RSC (right). **B.** Mean overlap of ensembles (left to right) between the final trial of each session in ACC (top) and RSC (bottom).

**Fig. S3.**
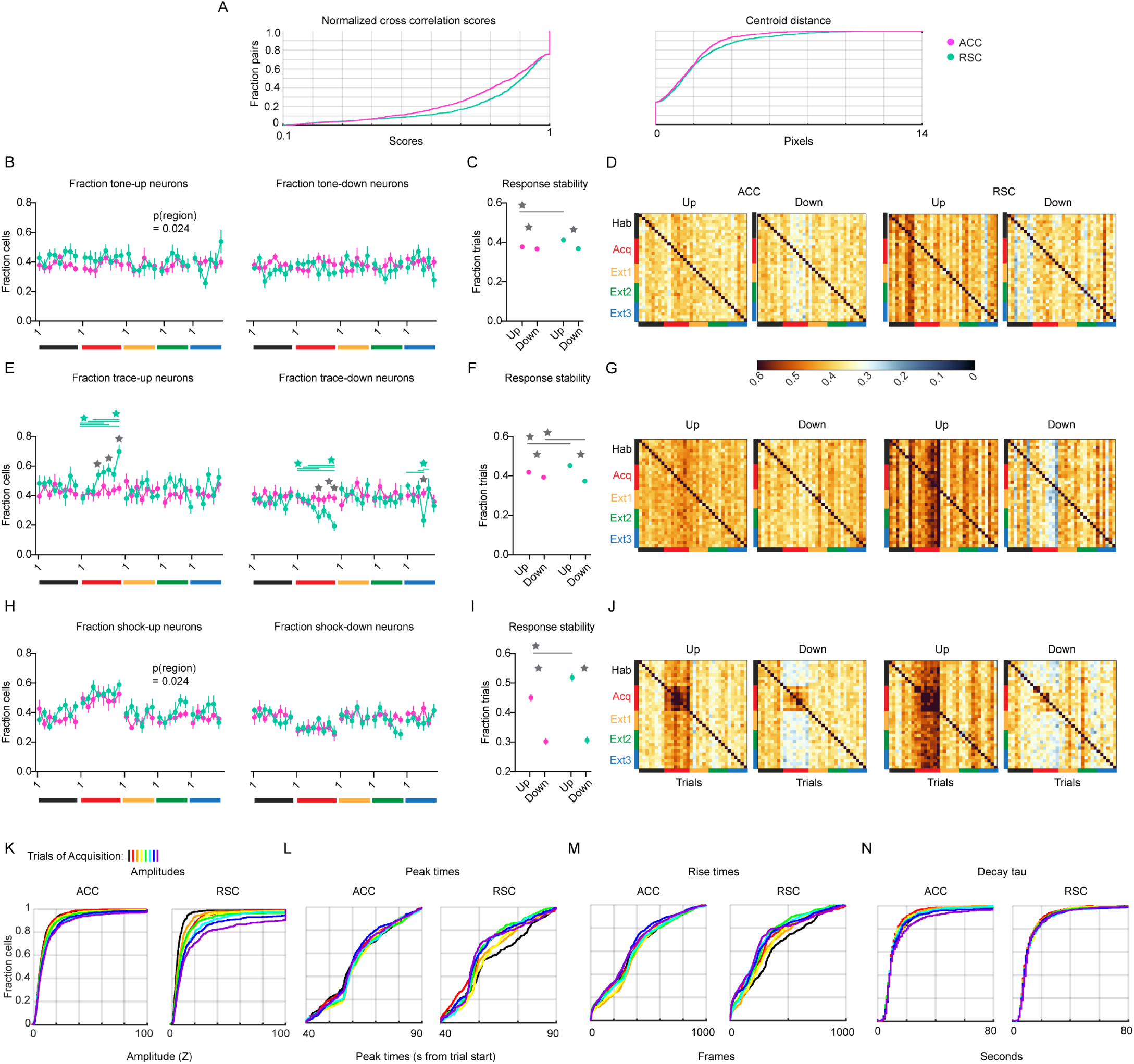
Neural tuning over TFC. **A.** Cumulative distributions of normalized cross-correlation scores (left) and centroid distances (right) between longitudinally registered ROIs in ACC (pink) and RSC (green). **B.** Fraction of significantly tone-upregulated (left; Two-Way RM ANOVA: pRegion_Ext1_ = 0.024, F(6,18) = 6.11) and -downregulated (right) neurons in each trial within each session. Colored bars indicate session ID. **C.** Fraction of trials that significantly modulated neurons maintained their tone-response properties (Two-Way ANOVA; pInteraction < 0.0001, F(1,1974) = 22.17). **D.** Heatmaps showing the overlap of significantly tone up- and down-regulated neurons in the ACC (left) and RSC (right) over trials. Colored bars indicate session ID. **E.** Fraction of significantly tone-upregulated (left; Two-Way RM ANOVA: pInteraction_Acq_ = 0.025, F(7,126) = 2.38; pTrial_Ext2_ = 0.032, F(5,90) = 2.58) and -downregulated (right; Two-Way RM ANOVA: pInteraction_Acq_ = 0.013, F(7,126) = 2.69; pInteraction_Ext3_ = 0.035, F(5,90) = 2.53) neurons in each trial within each session. Colored bars indicate session ID. **F.** Fraction of trials that significantly modulated neurons maintained their trace-response properties (Two-Way ANOVA; pInteraction < 0.0001, F(1,1974) = 44.65). **G.** Heatmaps showing the overlap of significantly trace up- and down-regulated neurons in the ACC (left) and RSC (right) over trials. Colored bars indicate session ID. **H.** Fraction of significantly shock-upregulated (left; Two-Way RM ANOVA: pRegion_Ext1_ = 0.024, F(1,18) = 6.11) and -downregulated (right) neurons in each trial within each session. Colored bars indicate session ID. **I.** Fraction of trials that significantly modulated neurons maintained their trace-response properties (Two-Way ANOVA; pInteraction = 0.0052, F(1,1974) = 7.84). **J.** Heatmaps showing the overlap of significantly trace up- and down-regulated neurons in the ACC (left) and RSC (right) over trials. Colored bars indicate session ID. **K.** Cumulative distributions of response amplitudes of ACC (top) and RSC (bottom) neurons over each trials, colored according to black-ROYGBIV from trials 1-8 of Acquisition. **L.** Same as E for peak times. **M.** Same as E for rise times. **N.** Same as E for decay times. Stars indicate p<0.05. Green stars indicate change in RSC over time; pink stars indicate change in ACC over time; gray stars with lines indicate change across regions over time; gray stars without lines indicate difference between regions. Statistics in Supplementary Table Rows 97-129.

**Supplementary Table.**
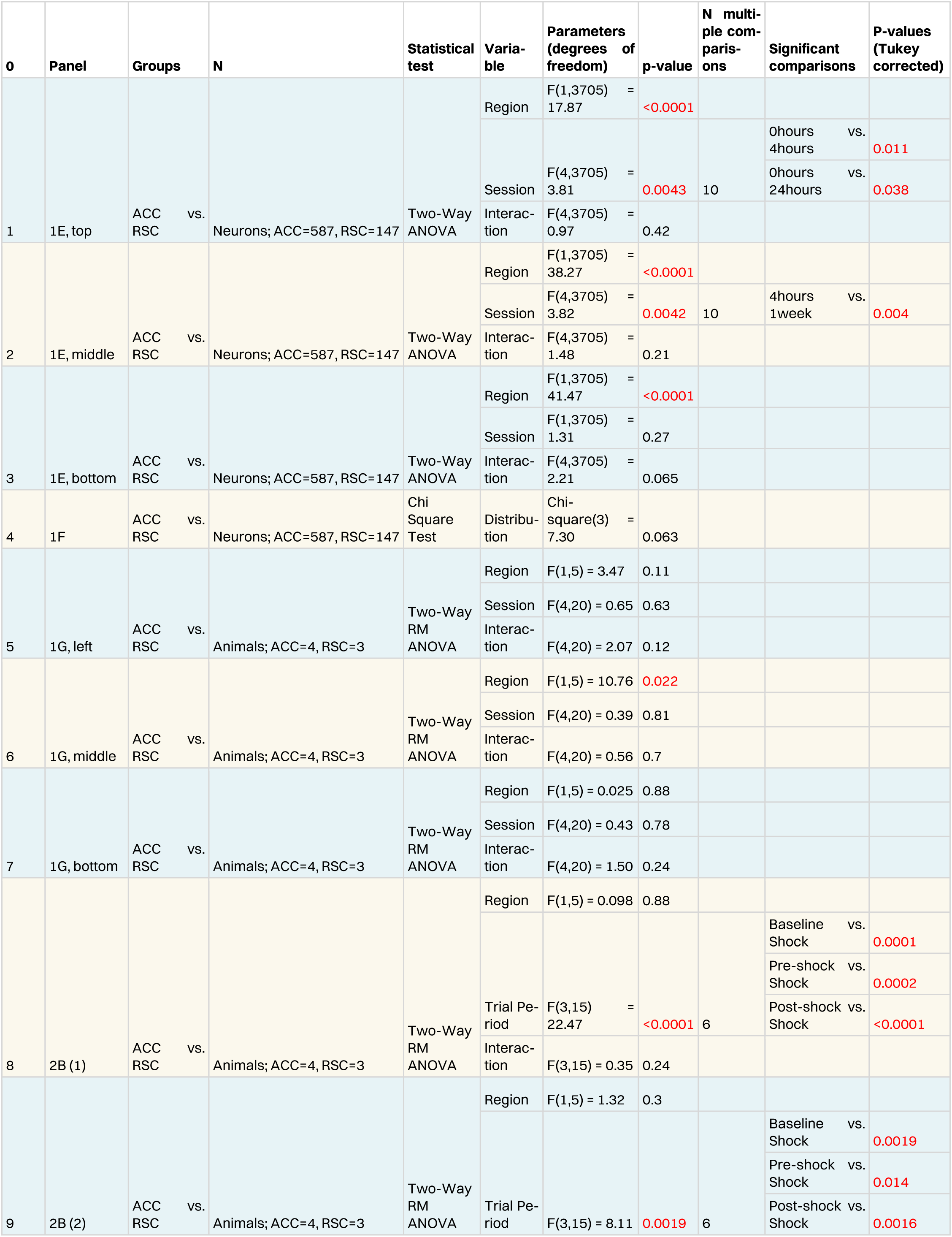
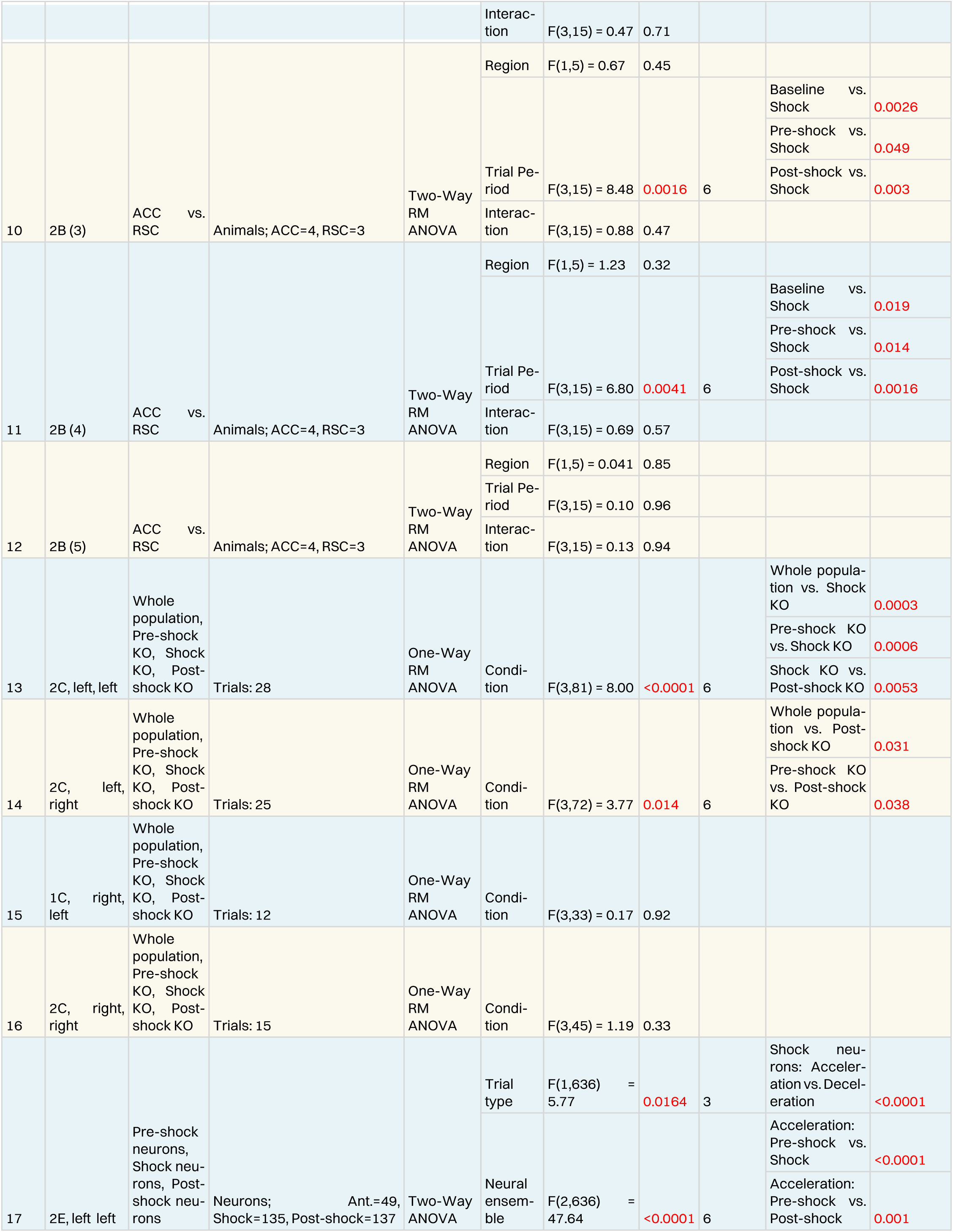
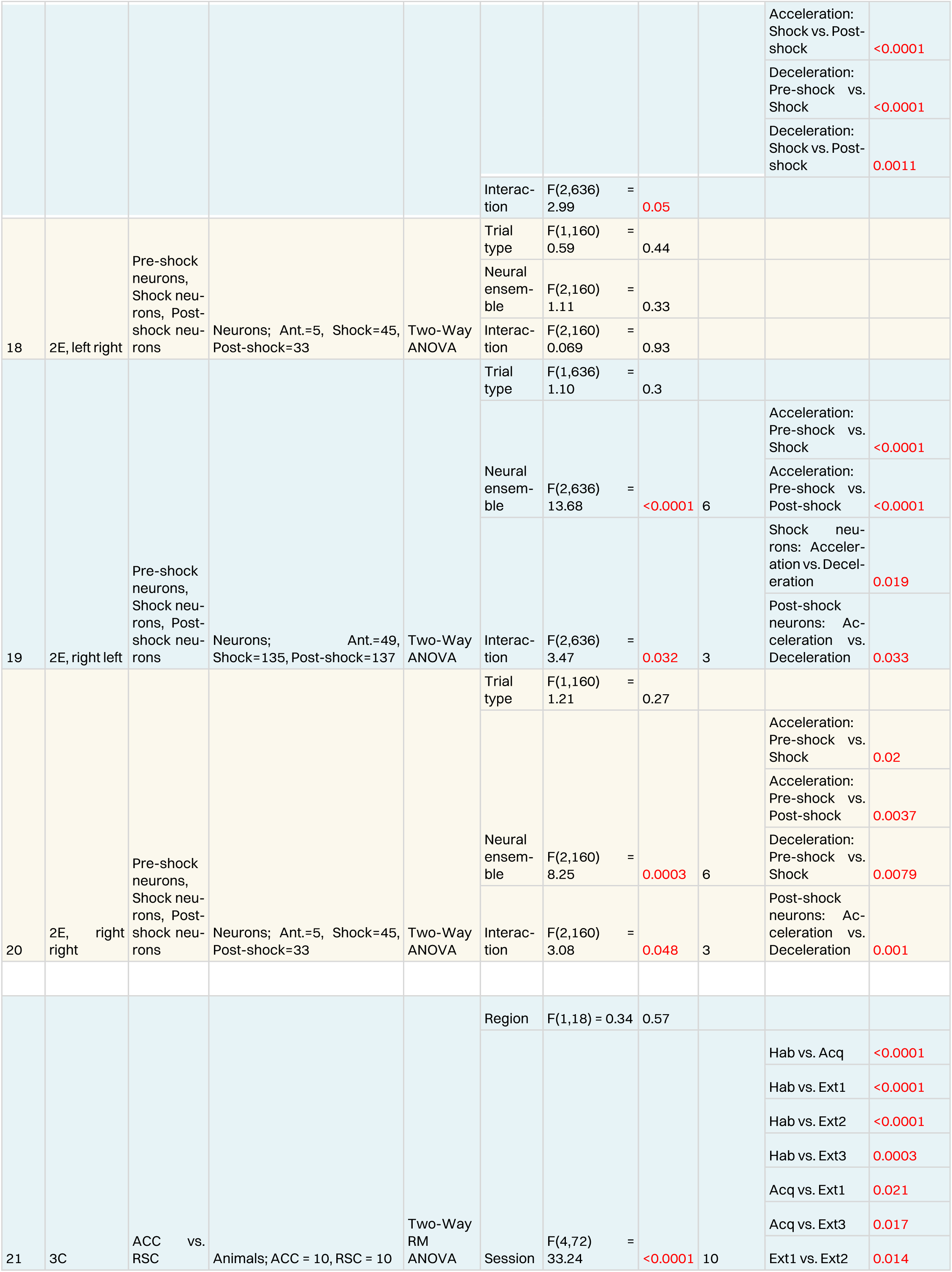
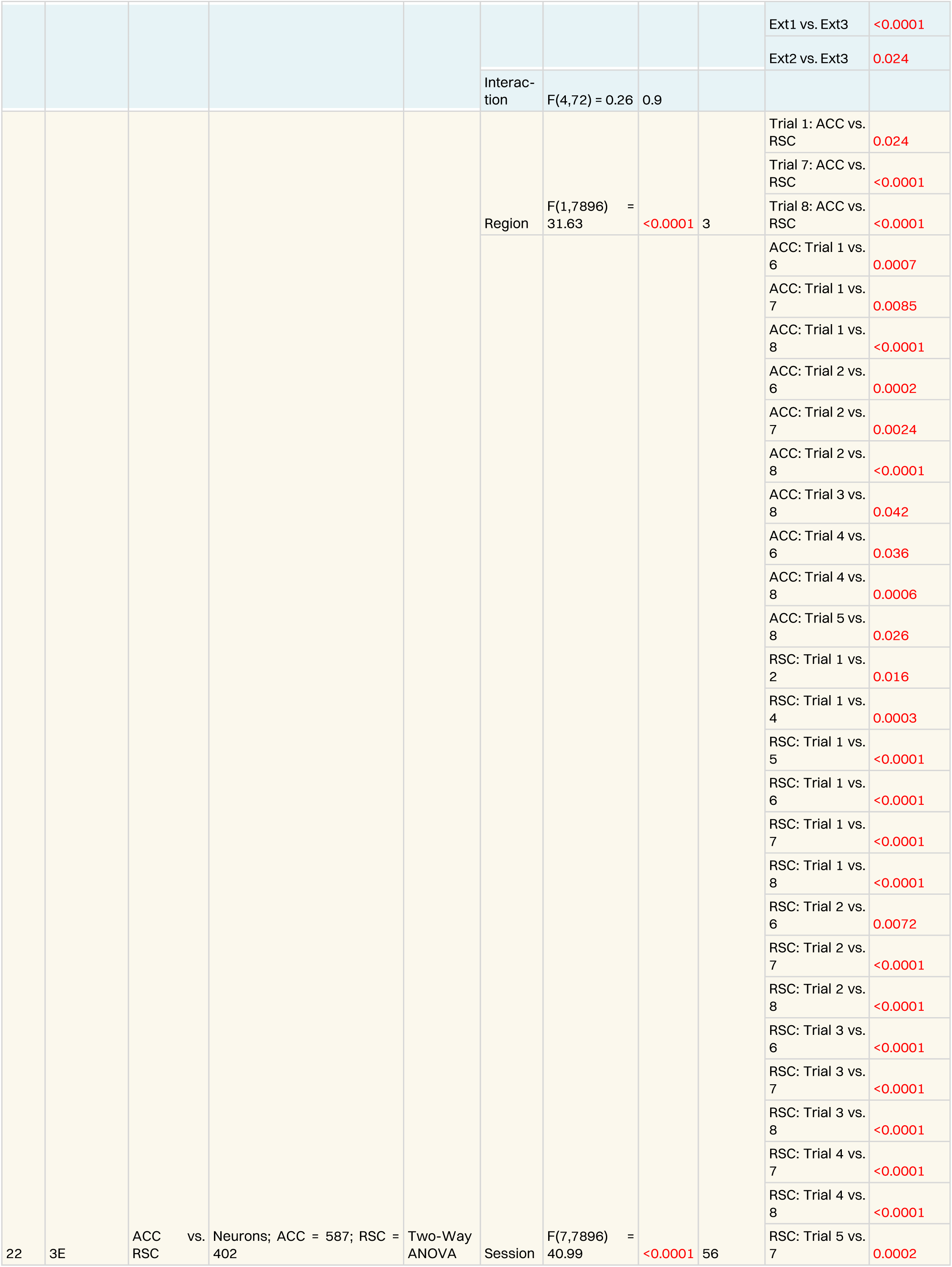
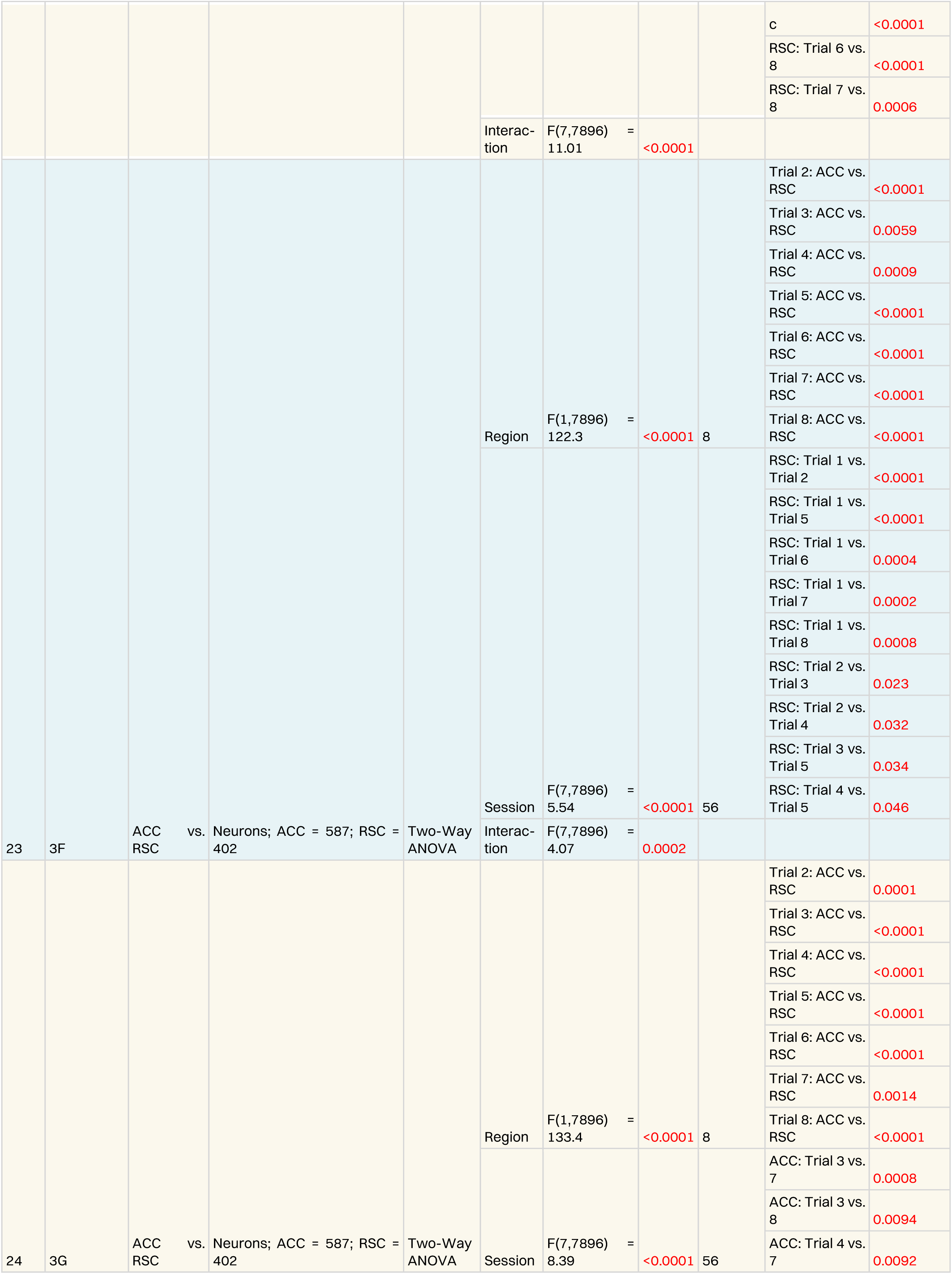
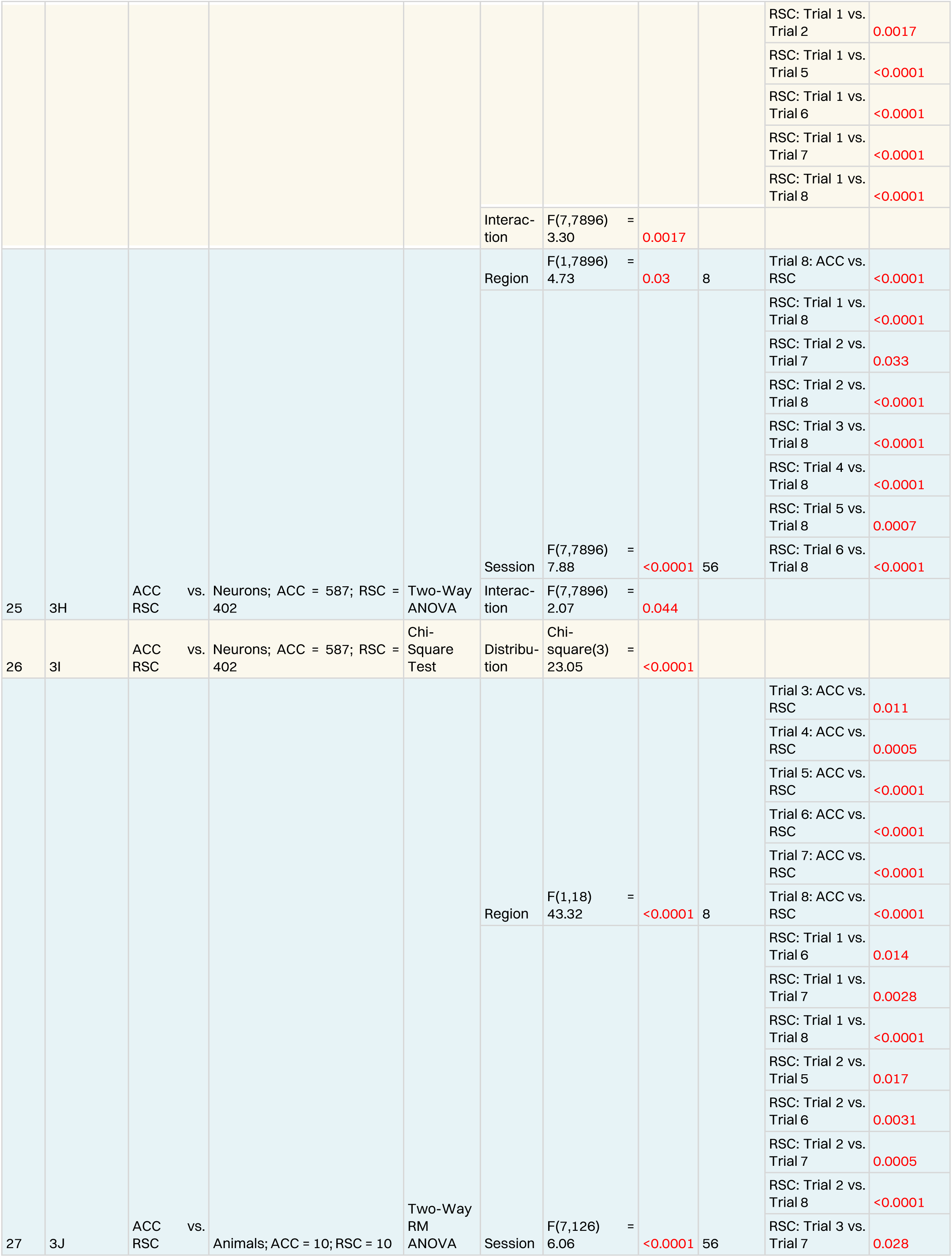
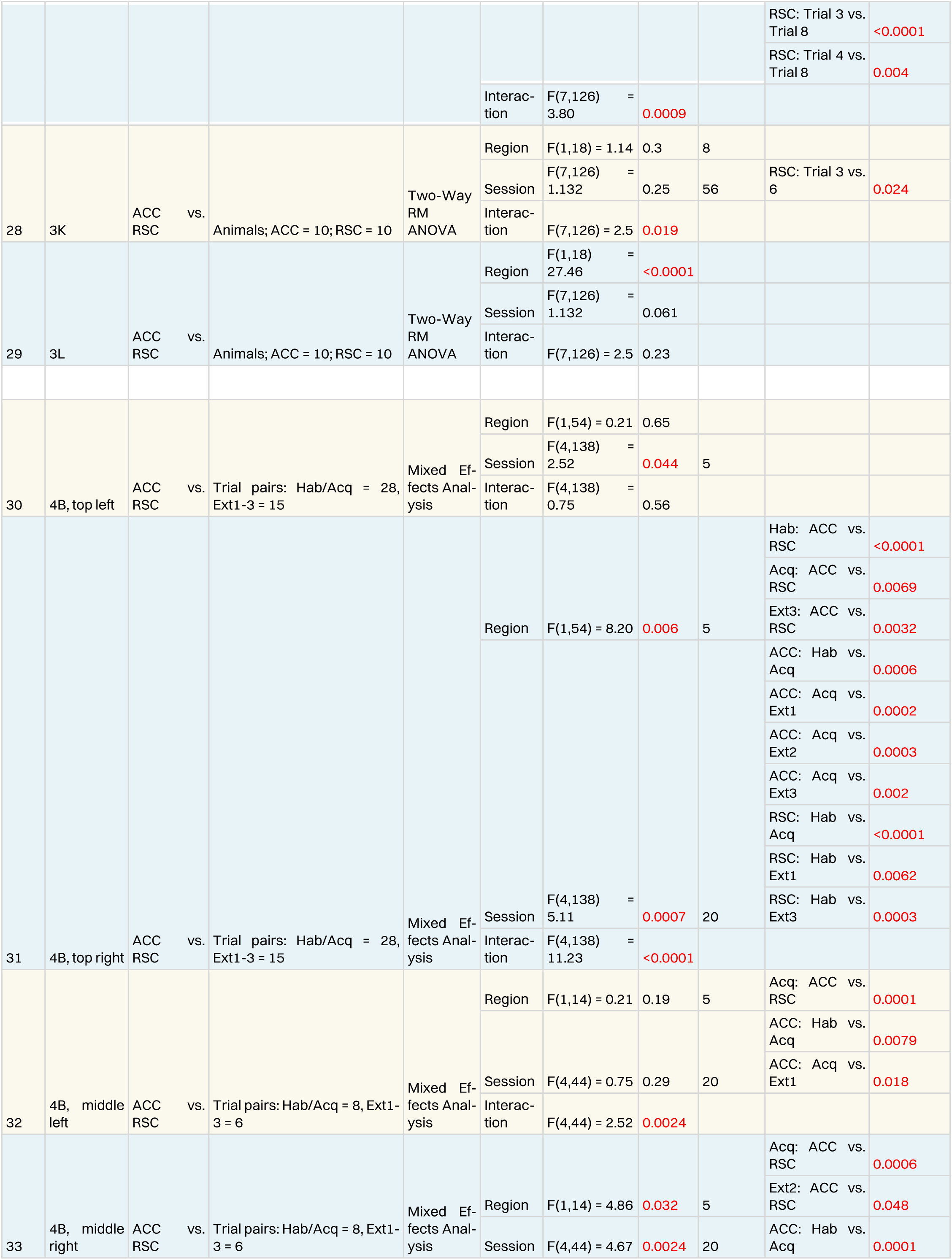
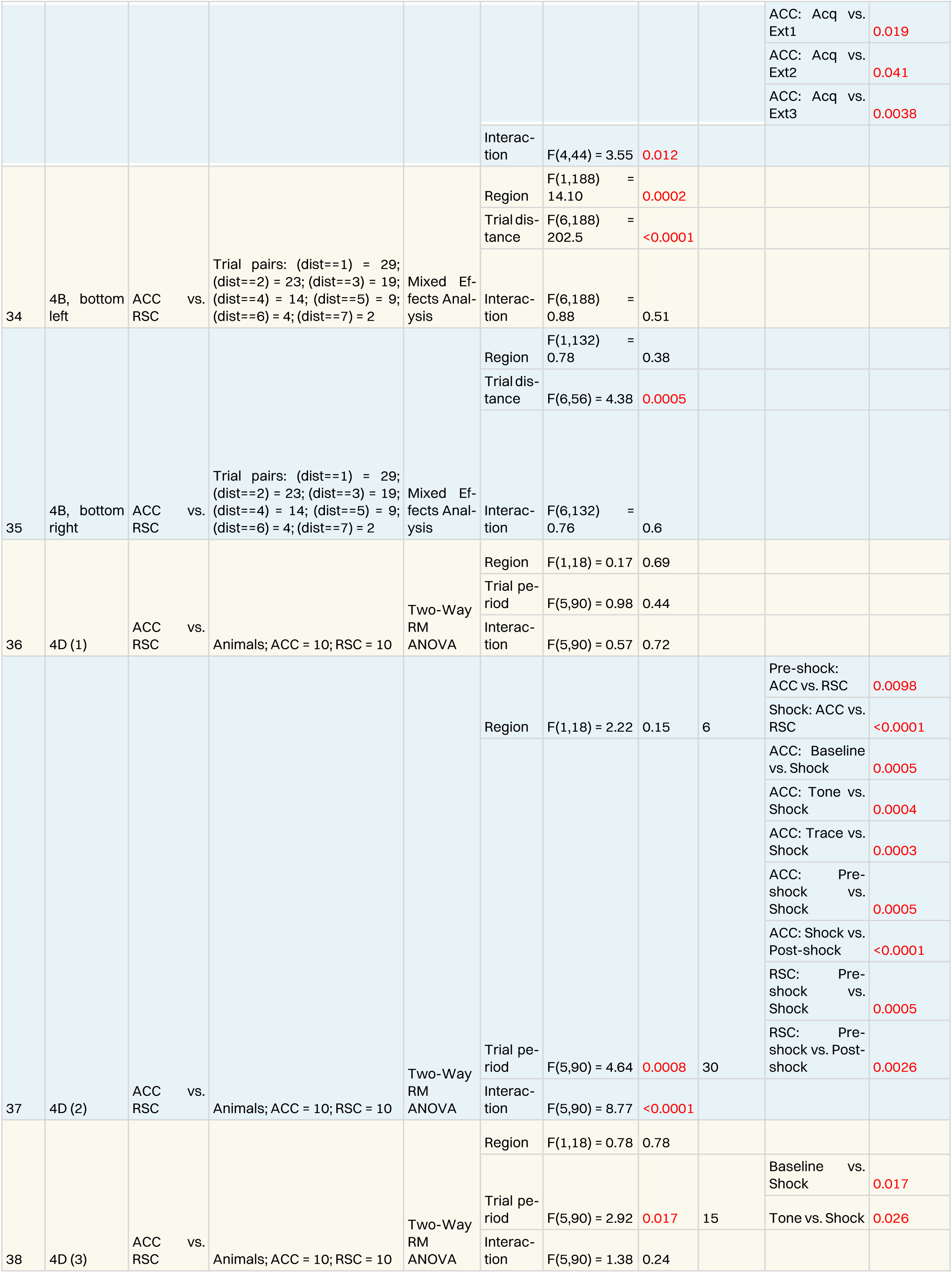
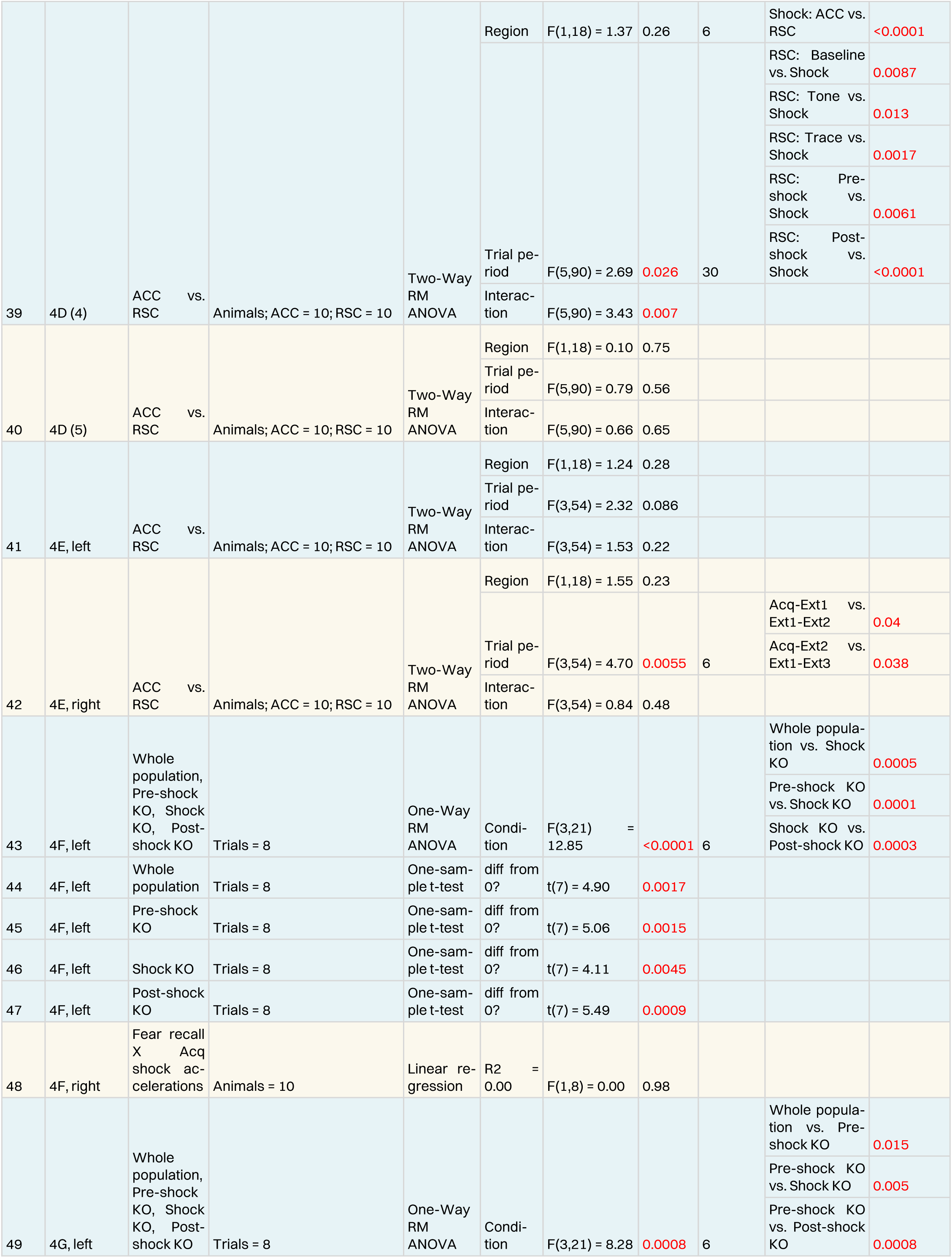
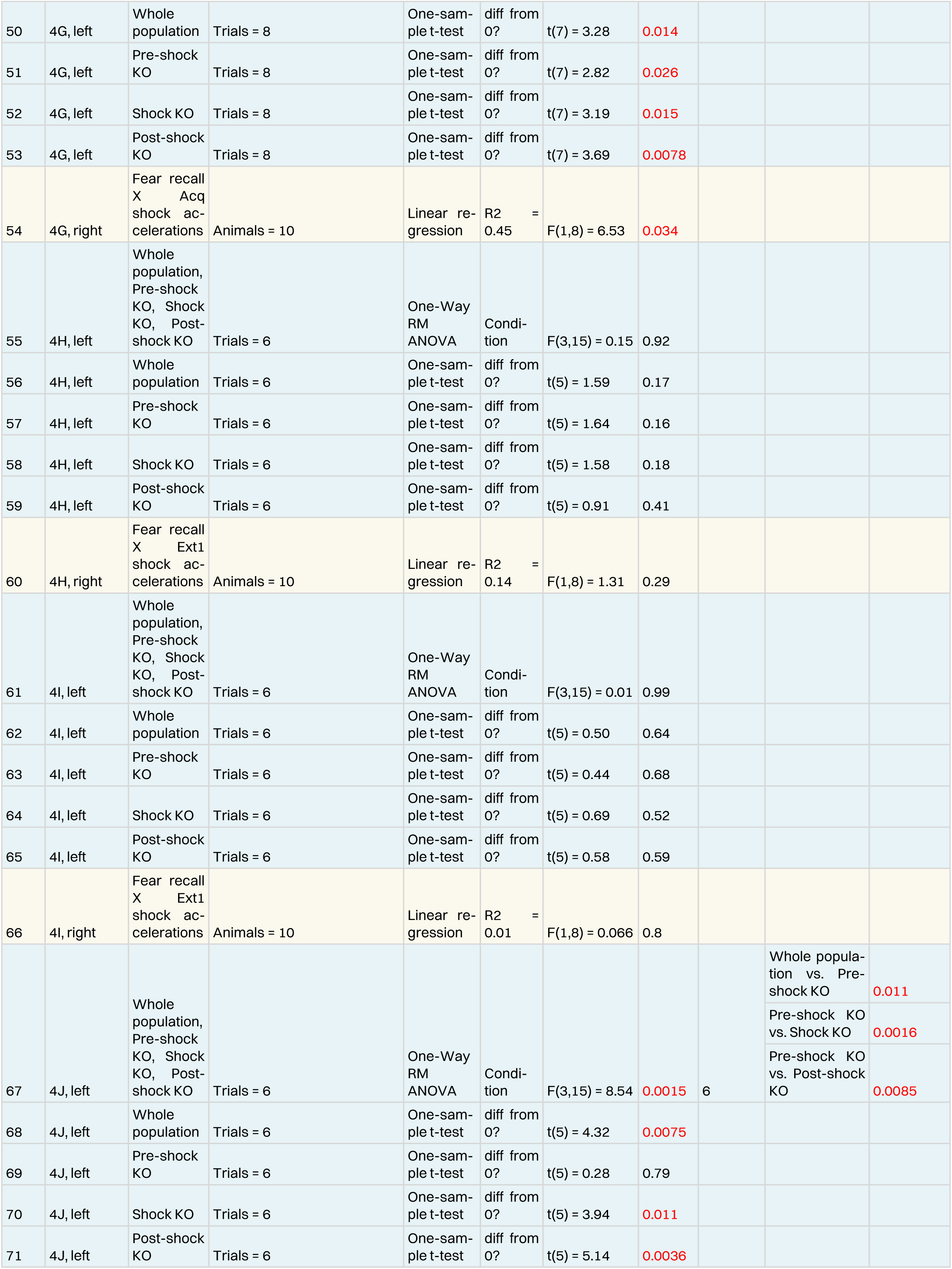
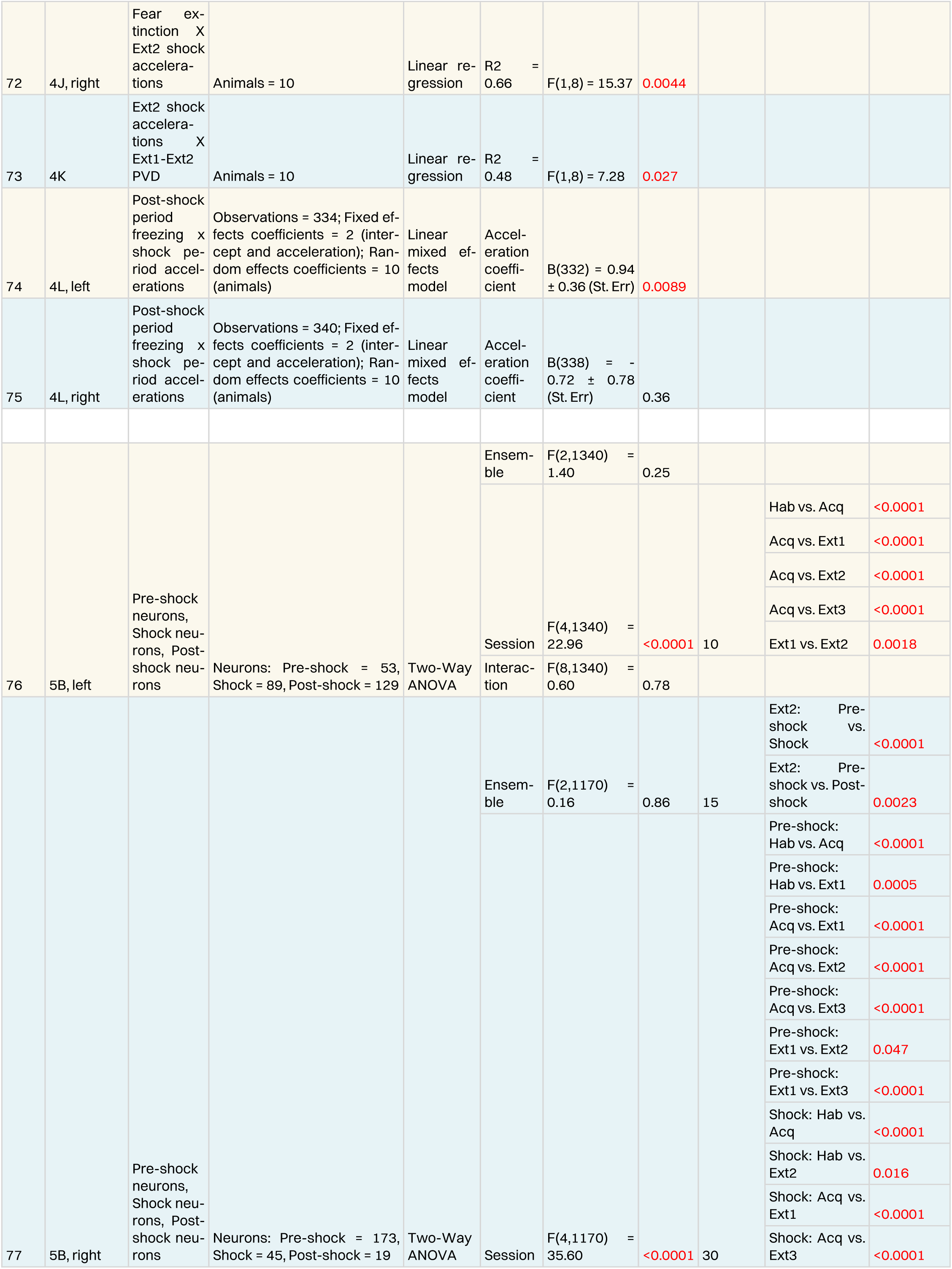
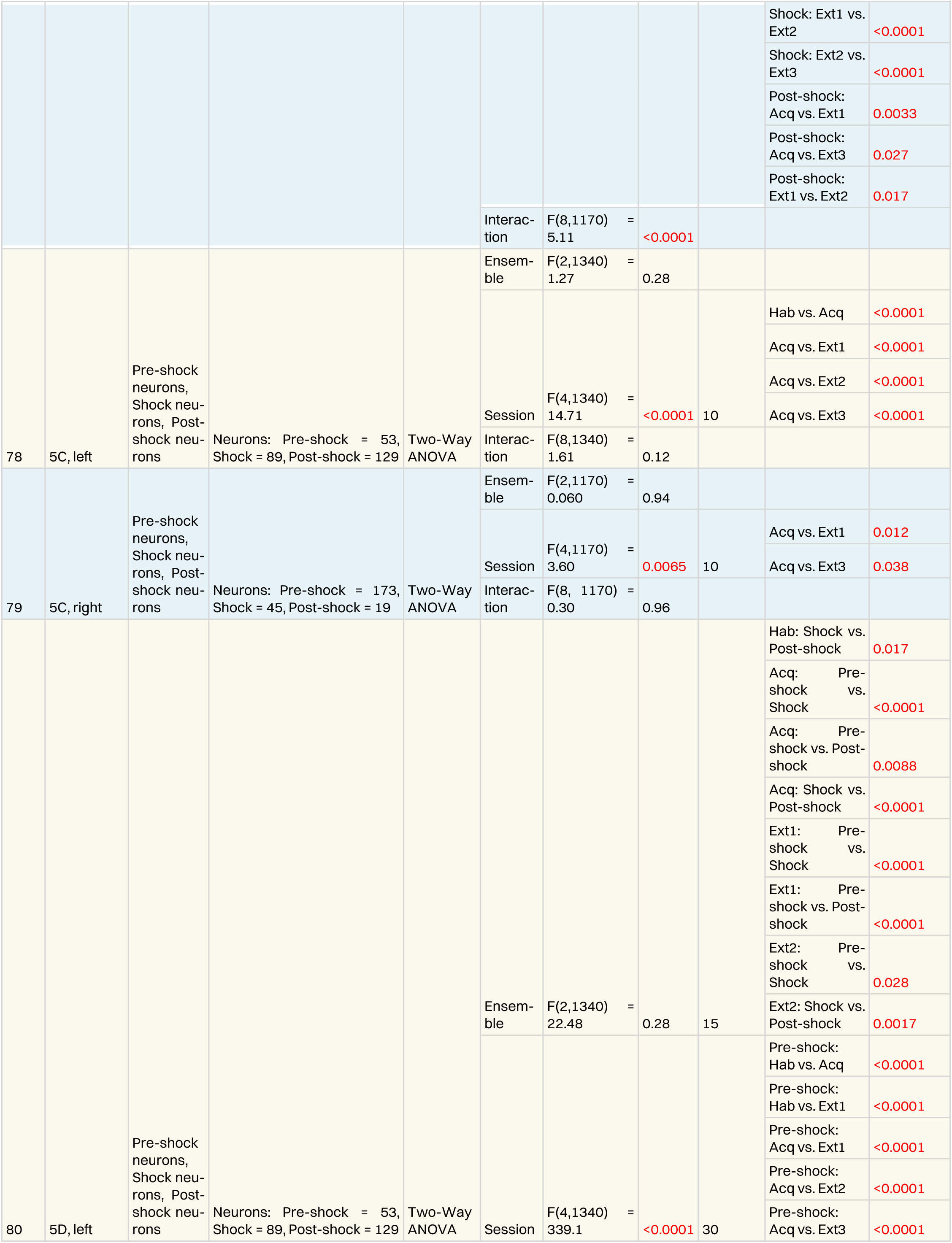
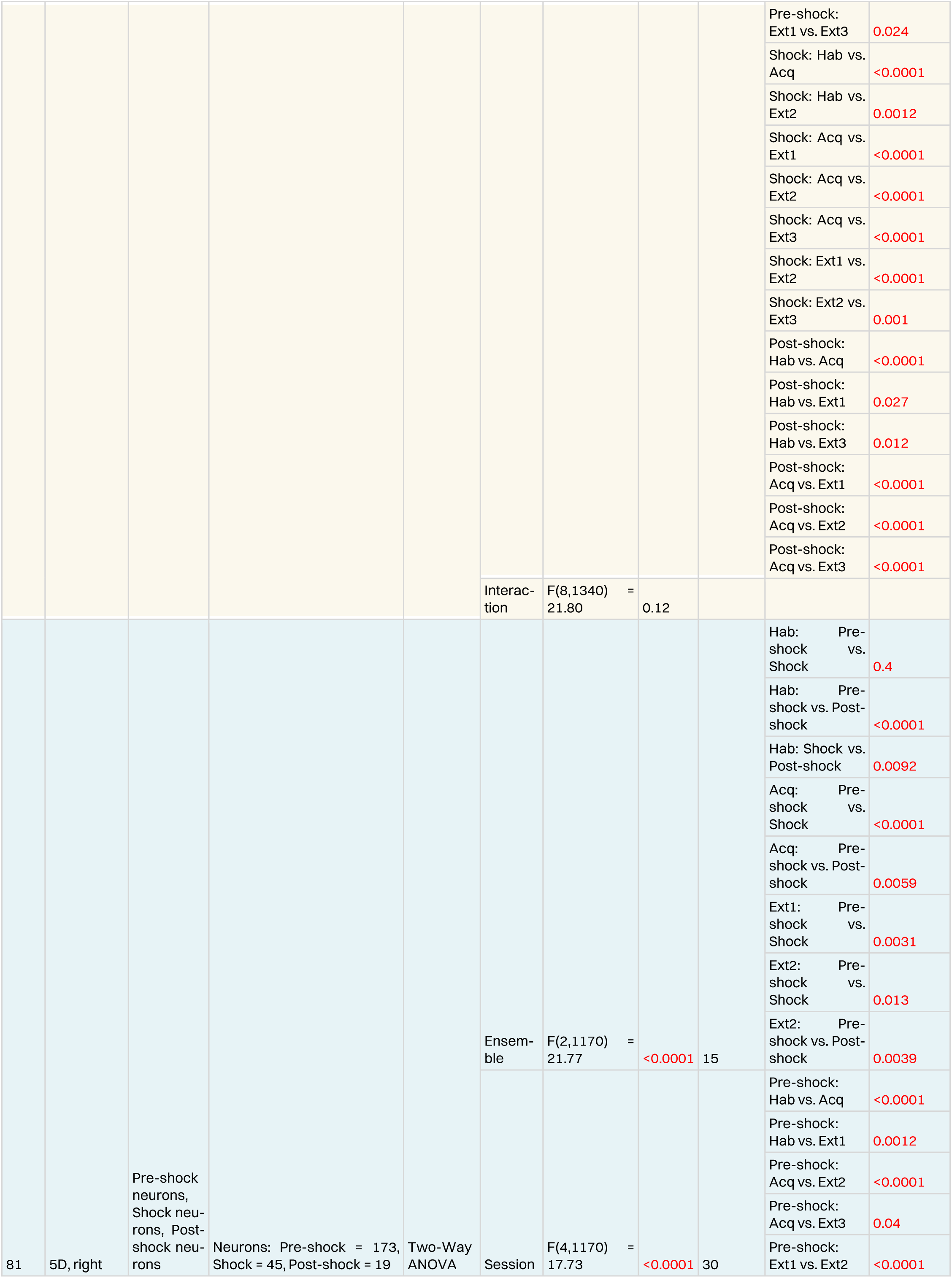
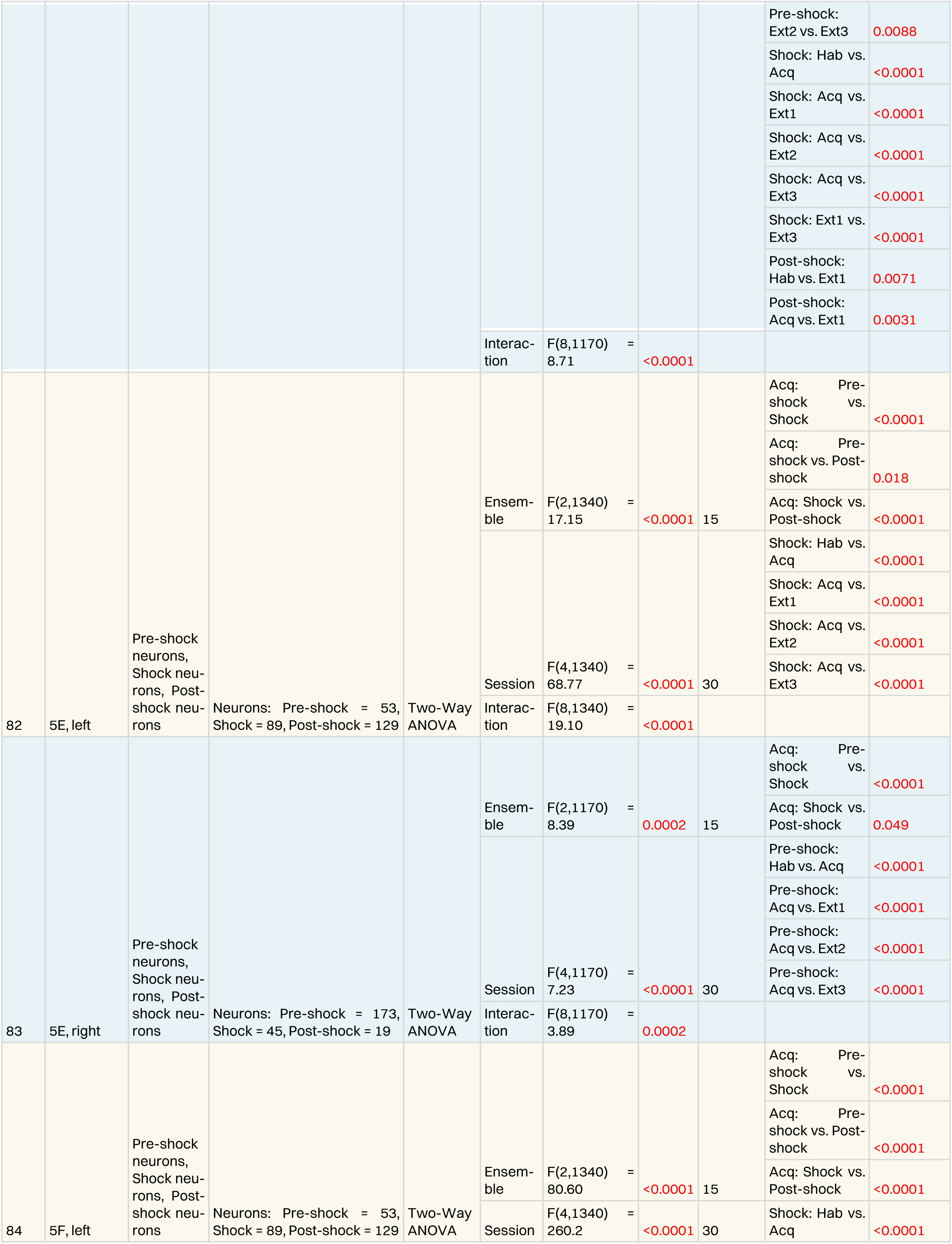
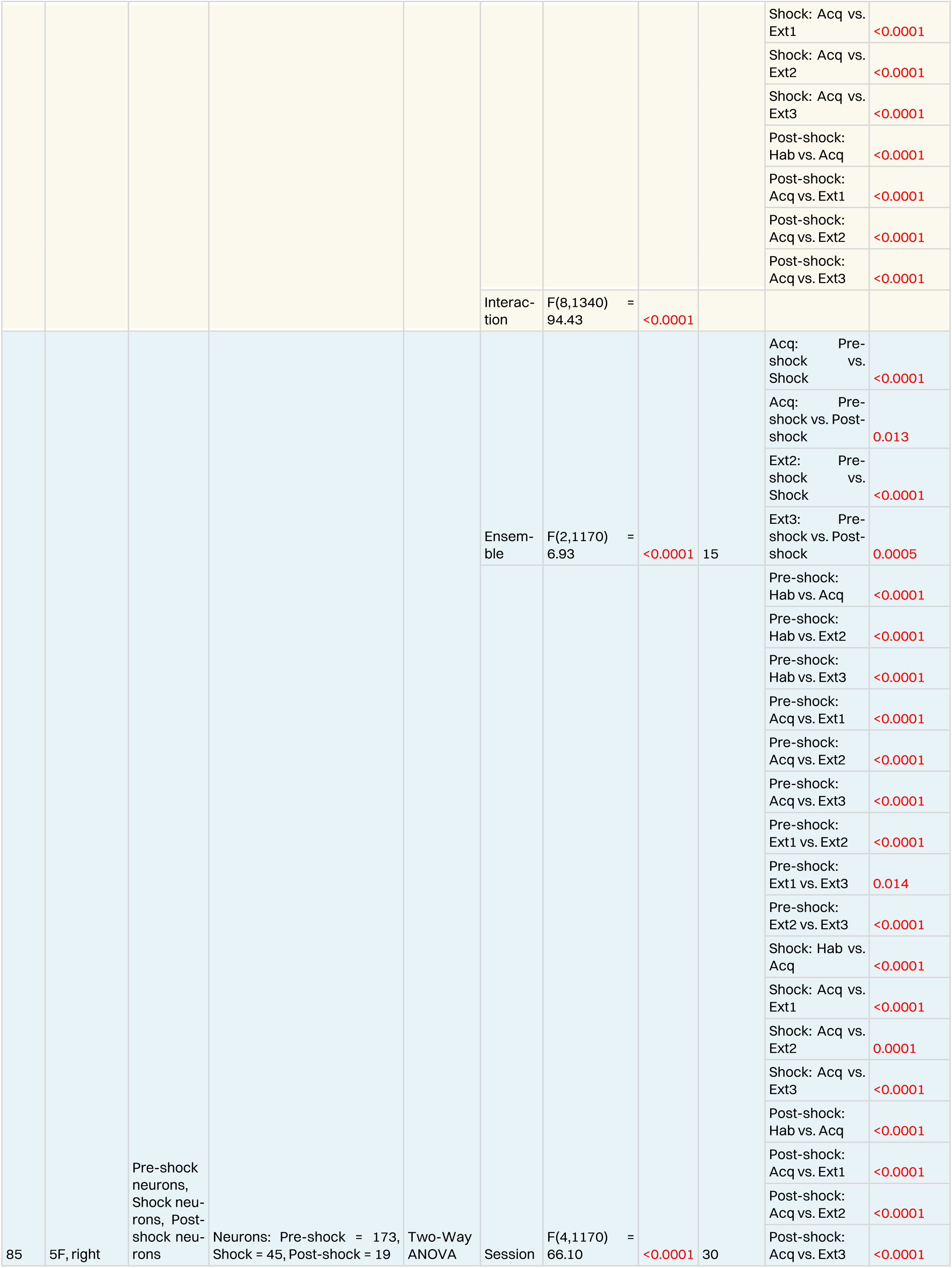
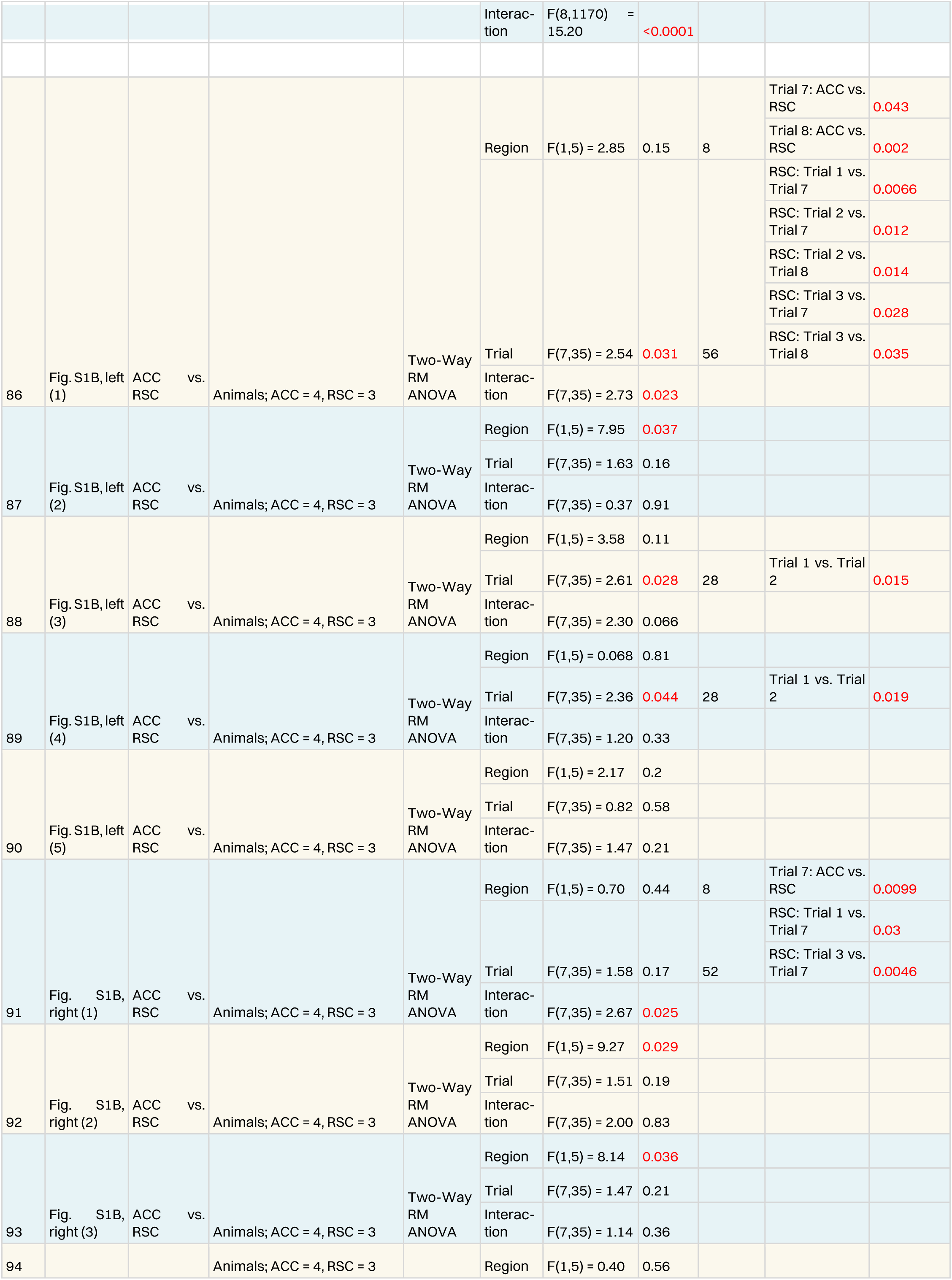
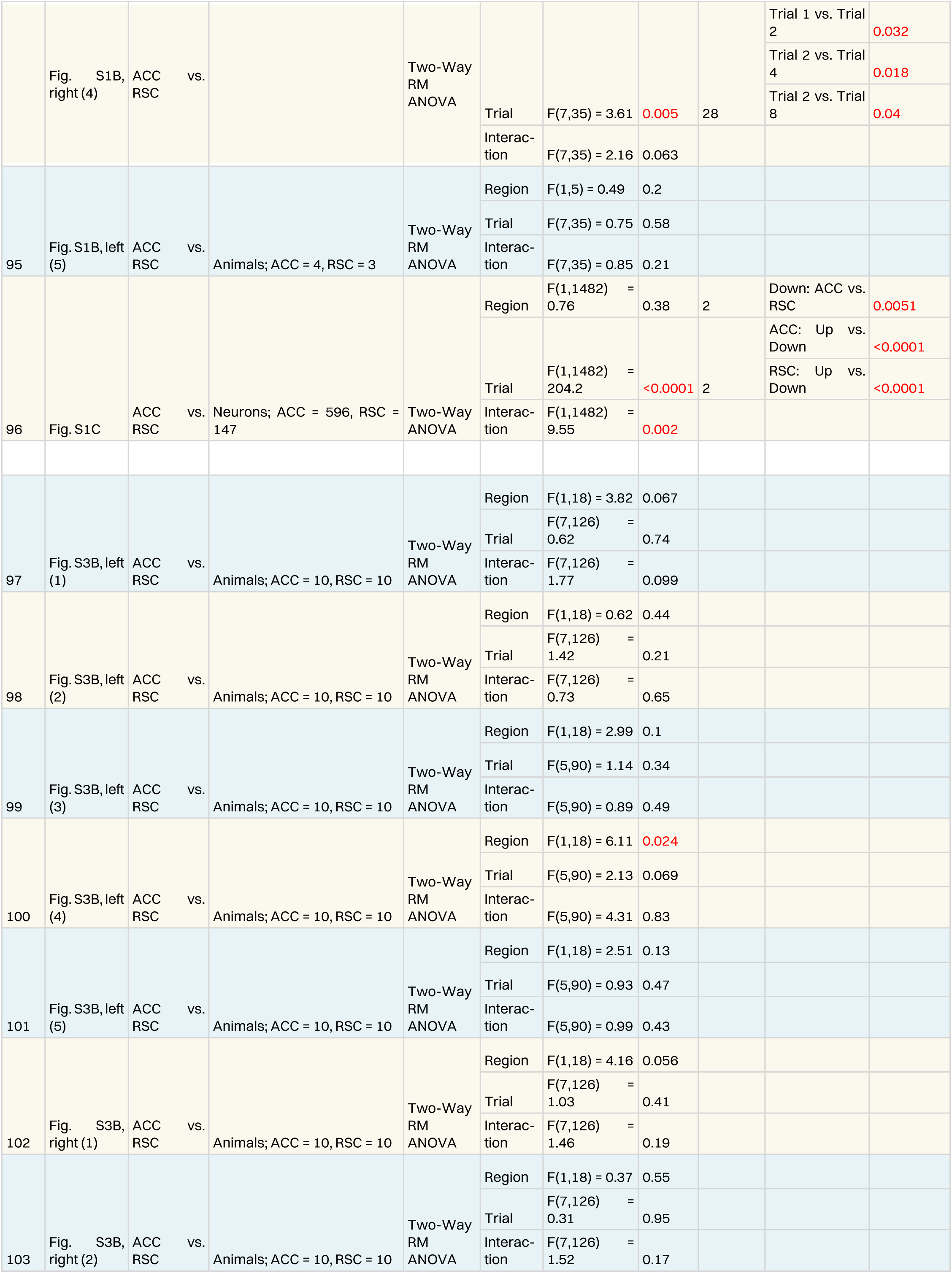
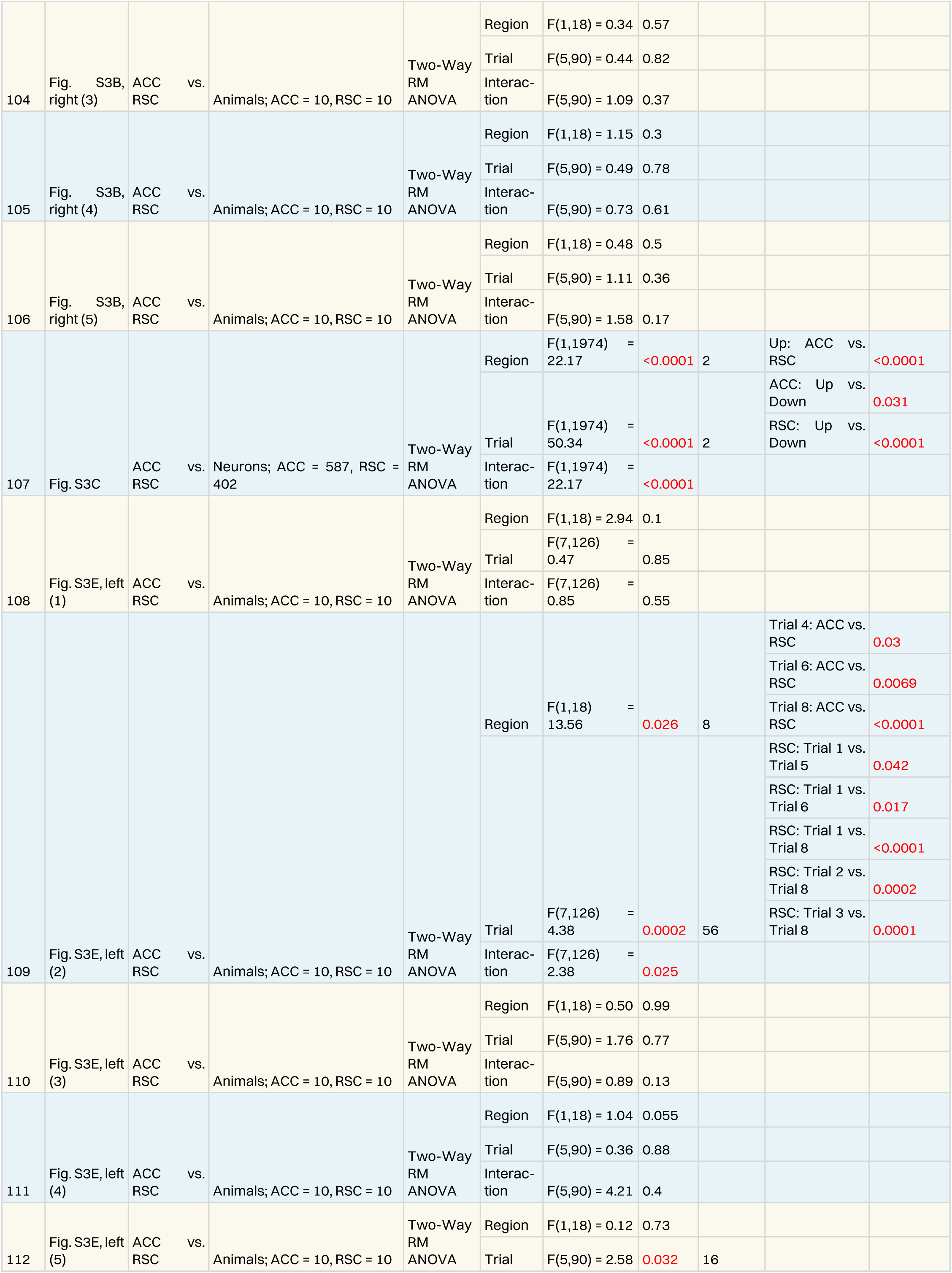
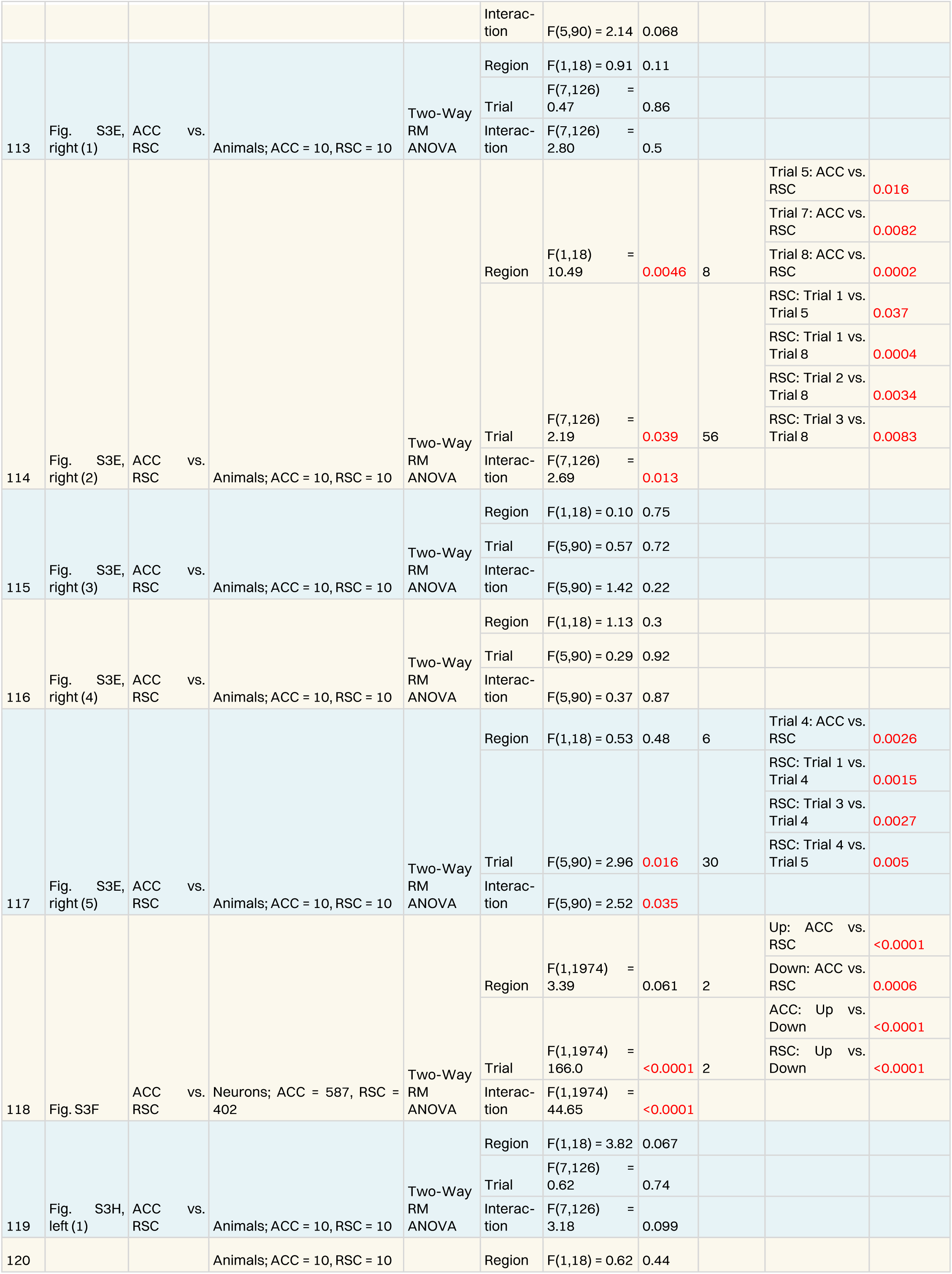
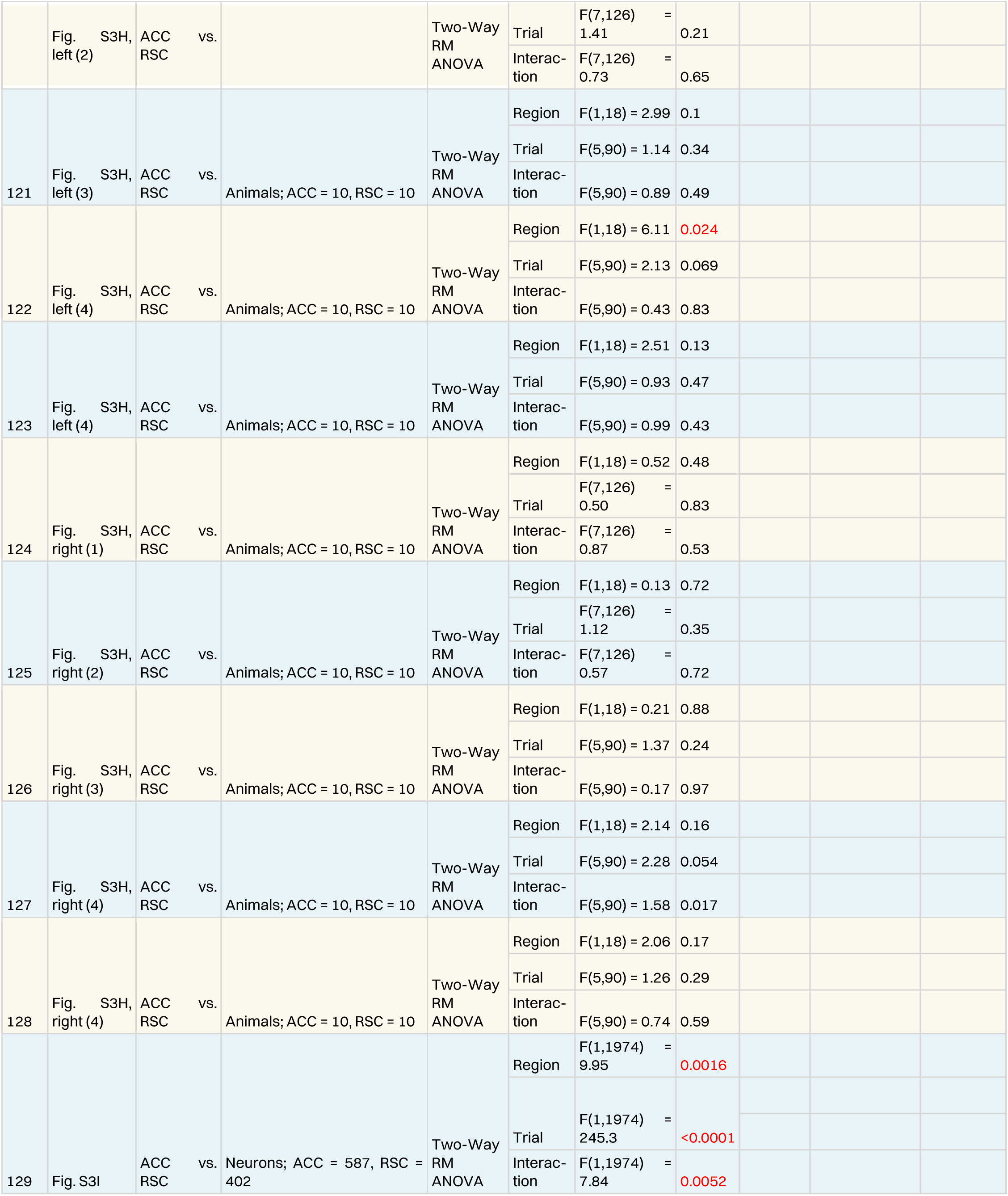
Statistical tests for all data presented.

